# Transcriptional silencing of *ALDH2* in acute myeloid leukemia confers a dependency on Fanconi anemia proteins

**DOI:** 10.1101/2020.10.23.352070

**Authors:** Zhaolin Yang, Yiliang Wei, Xiaoli S. Wu, Shruti V. Iyer, Moonjung Jung, Emmalee R. Adelman, Olaf Klingbeil, Melissa Kramer, Osama E. Demerdash, Kenneth Chang, Sara Goodwin, Emily Hodges, W. Richard McCombie, Maria E. Figueroa, Agata Smogorzewska, Christopher R. Vakoc

## Abstract

Hundreds of genes become aberrantly silenced in acute myeloid leukemia (AML), with most of these epigenetic changes being of unknown functional consequence. Here, we demonstrate how gene silencing can lead to an acquired dependency on the DNA repair machinery in AML. We make this observation by profiling the essentiality of the ubiquitin conjugation and ligation machinery in cancer cell lines using domain-focused CRISPR screening, which revealed Fanconi anemia (FA) proteins UBE2T (an E2) and FANCL (an E3) as unique dependencies in AML. We demonstrate that these dependencies are due to a synthetic lethal interaction between FA proteins and Aldehyde Dehydrogenase 2 (ALDH2), which function in parallel pathways to counteract the genotoxic effects of endogenous aldehydes. We provide evidence that DNA hypermethylation and transcriptional silencing of *ALDH2* occur in a recurrent manner in human AML patient samples, which is sufficient to confer FA pathway dependency in this disease. Taken together, our study suggests that targeting of the ubiquitination reaction catalyzed by FA proteins can eliminate ALDH2-deficient AML.

## INTRODUCTION

Organisms have evolved redundant genetic pathways for carrying out essential cellular processes. This often arises from gene duplication events, which produce paralogs that carry out overlapping functions (Kafri et al., 2009). Alternatively, non-homologous gene pairs can be redundant if they encode parallel pathways that regulate a shared cellular process (Innan and Kondrashov, 2010). For example, BRCA1 and PARP1 are biochemically distinct, yet function redundantly to repair DNA double stranded breaks (Bryant et al., 2005; Farmer et al., 2005). In this example of synthetic lethality, cells can tolerate a deficiency of either BRCA1 or PARP1 alone, yet a combined loss of these two factors causes cell death. While genetic redundancies support robustness during normal processes, cancer cells often lack such redundancies owing to genetic or epigenetic alterations (Cereda et al., 2016; O’Connor, 2015). This offers therapeutic opportunities, such as drugs that block PARP1 in cancers caused by a genetic deficiency of *BRCA1* (Bryant et al., 2005; Farmer et al., 2005). Due to its therapeutic significance, the identification of synthetic lethal genetic interactions remains an important objective in the study of human cancer (Kaelin, 2005).

In recent years, there has been a resurgent interest in the ubiquitination machinery as targets for cancer therapy. Ubiquitin is a 8.6 kDa protein that is covalently attached to lysine side chains as a form of post-translational regulation of protein stability and function (Komander and Rape, 2012). The cascade reaction of ubiquitination requires the consecutive action of three enzymes: a ubiquitin-activating enzyme (E1), ubiquitin-conjugating enzymes (E2), and ubiquitin ligases (E3) (Grabbe et al., 2011). With ~40 E2 and over 600 putative E3 proteins encoded in the human genome, the ubiquitination machinery regulates many aspects of cell biology, including the cell cycle, DNA repair, and transcription (Senft et al., 2018). In addition, it has been established that the function of E2 and E3 proteins can be modulated with small molecules (Hoeller and Dikic, 2009; Skaar et al., 2014). For example, the E3 ligase protein MDM2 can be inhibited with small molecules to stabilize p53 and promote apoptosis of cancer cells (Hatakeyama, 2011). In addition, small molecules have been identified that alter the specificity of E3 ligases to trigger proteasome-mediated degradation of neo-substrates (Kronke et al., 2014; Lu et al., 2014). Despite the clear therapeutic potential of the ubiquitination machinery, there have been few efforts to date aimed at identifying E2/E3 dependencies in cancer cells using genetic screens.

The Fanconi anemia (FA) pathway, comprised of more than 20 protein components, repairs DNA interstrand crosslinks (ICLs) (Ceccaldi et al., 2016). A key step in the activation of the FA repair pathway is the mono-ubiquitination of FANCD2 and FANCI, which is performed by UBE2T/FANCT (an E2 conjugating enzyme) and FANCL (an E3 ligase) (Ceccaldi et al., 2016; Kottemann and Smogorzewska, 2013). Once ubiquitinated, a FANCD2/FANCI heterodimer encircles the DNA (Alcón et al., 2020; Wang et al., 2020), which facilitates lesion processing by nucleases and completion of the repair by homologous recombination (Ceccaldi et al., 2016; Wang and Smogorzewska, 2015). While DNA crosslinks can be caused by exogenous mutagens, emerging evidence suggests that endogenously produced aldehydes are an important source of this form of DNA damage (Langevin et al., 2011). Loss-of-function mutations in FA genes lead to Fanconi anemia, a genetic disorder characterized by bone marrow failure and a predisposition to leukemia and aerodigestive tract squamous cell carcinoma (Kottemann and Smogorzewska, 2013). While a deficiency in FA proteins is cancer-promoting in certain tissue contexts, for the majority of human cancers the FA pathway remains intact and therefore may perform an essential function in established cancers. To our knowledge, a role of FA proteins as cancer dependencies has yet to be identified.

Here, we discovered an acquired dependency on FA proteins in a subset of AML that lacks expression of Aldehyde Dehydrogenase 2 (ALDH2). We show that epigenetic silencing of *ALDH2* occurs in a recurrent manner in human AML and is sufficient to confer FA dependency in this disease. Our study suggests that blocking the UBE2T/FANCL-mediated ubiquitination can selectively eliminate ALDH2-deficient AML cells, but would spare ALDH2-expressing normal cells present in the majority of human tissues.

## METHODS

### Cell Lines

Leukemia lines MOLM-13, NOMO-1, MV4-11, ML-2, HEL, SET-2, THP-1, U937, K562, pancreatic ductal adenocarcinoma (PDAC) lines AsPC-1, CFPAC-1, SUIT-2, PANC-1, MIAPaCa-2, small cell lung cancer (SCLC) lines NCI-H211, NCI-H82, DMS114, murine RN2c (MLL-AF9/Nras^G12D^ AML) were cultured in RPMI supplemented with 10% FBS. SEM cells were cultured in IMDM with 10% FBS. OCI-AML3 cells were cultured in alpha-MEM with 20% FBS. KASUMI-1 cells were cultured in RPMI supplemented with 20% FBS. NCI-H1048 cells were cultured in DMEM:F12 supplemented with 0.005 mg/mL insulin, 0.01 mg/mL transferrin, 30 nM sodium selenite, 10 nM hydrocortisone, 10 nM β-estradiol, 4.5 mM L-glutamine and 5% FBS. RH30, RD, RH4, CTR (rhabdomyosarcoma), HEK293T were cultured in DMEM supplemented with 10% FBS and 4.5 mM L-glutamine. MLL-AF9 (MA9, engineered human AML) cells were cultured in IMDM supplemented with 20% FBS, 10 ng/mL SCF, 10 ng/mL TPO, 10 ng/mL FLT3L, 10 ng/mL IL-3, 10 ng/mL IL-6. MLL-AF9/Nas^G12D^, MLL-AF9/FLT3^ITD^ (engineered human AML) cells were cultured in IMDM with 20% FBS. Murine NIH-3T3 cells were cultured in DMEM with 10% bovine calf serum. Penicillin/Streptomycin was added to all media. All cell lines were cultured at 37 °C with 5% CO_2_, and were confirmed mycoplasma negative.

### Construction of E2 and E3 domain-focused sgRNA library

The list of E2 and E3 genes was retrieved from HUGO Gene Nomenclature Committee (HGNC) resource. The E2 and E3 enzymatic functional domain annotation was retrieved from the NCBI Conserved Domains Database. Six to ten independent sgRNAs were designed against each individual domain based while minimizing off-target effects (Hsu et al., 2013). The domain targeting and spike-in positive/negative control sgRNA oligonucleotides were synthesized using an array platform (Twist Bioscience) and then amplified by PCR. The PCR products were cloned into the BsmBI-digested optimized sgRNA lentiviral expression vector LRG2.1T using a Gibson assembly kit (New England Biolabs; Cat. No. E2611). The LRG2.1T vector was derived from a lentiviral U6-sgRNA-EFS-GFP expression vector (Addgene: #65656). The pooled plasmids library was subjected to deep-sequencing analysis on a MiSeq instrument (Illumina) to verify the identity and representative of sgRNAs in the library. It was confirmed that 100% of the designed sgRNAs were cloned in the LRG2.1T vector and that the abundance of >95% of individual sgRNA constructs was within 5-fold of the mean.

### Virus production and transduction

Lentivirus was produced in HEK293T cells by transfecting plasmids together with helper packaging plasmids (VSVG and psPAX2) using polyethylenimine (PEI 25000; Polysciences; Cat. No. 23966-1) transfection reagent. HEK293T cells were plated in 10 cm culture dishes and were transfected when confluency reached ~80-90%. Five plates of HEK293T were used to ensure the representation of library. For one 10 cm dish of HEK293T cells, 10 μg of plasmid DNA, 5 μg of pVSVG and 7.5 μg psPAX2 and 64 μL of 1 mg/mL PEI were mixed, incubated at room temperature for 20 min and then added to the cells. The media was changed to fresh media 6-8 h post-transfection. The media containing lentivirus was collected at 24h, 48h and 72h post transfection and pooled together. Virus was filtered through 0.45 μM non-pyrogenic filter.

For shRNA knock-down experiments, retrovirus was produced in Plat-E cells, which were transfected with retroviral DNA (MLS-E plasmid), VSVG, and Eco helper plasmids in a ratio of 10:1:1.5. The media was changed to fresh media 6-8 h post-transfection. Retrovirus-containing supernatant was collected at 24, 48 and 72 h after transfection and pooled together. Virus was filtered through 0.45 μM non-pyrogenic filter.

For both lentivirus and retrovirus infection, target cells were mixed with corresponding volume of virus supplemented with 4 μg/mL polybrene, and then centrifuged at 600 x g for 40 min at room temperature. If selection was needed for stable cell line establishment, corresponding antibiotics (1 μg/mL puromycin, 1 mg/mL G418) were added 72 h post infection.

### Plasmid construction, sgRNA and shRNA cloning

For CRISPR screening, the optimized sgRNA lentiviral expression vector (LRG2.1T) and the lentiviral human codon-optimized *Streptococcus pyogenes* Cas9 vector (LentiV_Cas9_Puro, Addgene: 108100) were used. For the competition-based proliferation assays, sgRNAs were cloned into the LRG2.1T vector using BsmBI restriction site. LRCherry2.1 was derived from LRG2.1T by replacing GFP with mCherry CDS. All sgRNA and shRNA sequences used in the study are listed in Supplementary Table 2. For the cDNA rescue experiments, the cDNA of UBE2T, FANCL, ALDH1A1, or ALDH2 were cloned into the LentiV_Neo vector using the In-Fusion cloning system (Takara Bio; Cat. No. 121416). The CRISPR-resistant synonymous mutant of UBE2T and FANCL, and catalytic mutant of UBE2T and ALDH2 were cloned using PCR mutagenesis. shRNAs targeting FANCD2 and CDK1 were cloned into the mirE-based retroviral shRNA expression vector MLS-E.

### Pooled negative-selection CRISPR screening and data analysis

CRISPR-based negative selection screening was performed in Cas9-expressing cancer cell lines, which were established by infection with LentiV-Cas9-Puro vector and selected with puromycin. The lentivirus of the pooled sgRNA library targeting the functional domains of ubiquitination genes was produced as described above. Multiplicity of infection (MOI) was set to 0.3-0.4 to ensure a single sgRNA transduction per cell. To maintain the representation of sgRNAs during the screen, the number of sgRNA infected cells was kept to 800 times the number of sgRNAs in the library. Cells were harvested at three days post-infection as an initial reference time point. Cells were cultured for 12 population doublings and harvested for the final time point. Genomic DNA was extracted using QIAamp DNA mini kit (QIAGEN; Cat. No. 51304) according to the manufacture’s instruction.

The extracted genomic DNA was used for sequencing library preparation. Briefly, the sgRNA cassette was PCR amplified from genomic DNA (~200 bp) using high-fidelity polymerase (Phusion master mix, Thermo Fisher; Cat. No. F531S). The PCR product was end-repaired with T4 DNA polymerase (NEB; Cat. No. B02025), DNA Polymerase I, Large (Klenow) fragment (NEB; Cat. No. M0210L) and T4 polynucleotide kinase (NEB; Cat. No. M0201L). A 3’ A overhang was added to the ends of the blunted DNA fragment with Klenow Fragment (3’-5’ exo; NEB; Cat. No. M0212L). The DNA fragments was then ligated with diversity-increased custom barcodes (Shi et al., 2015), with Quick ligation kit (NEB; Cat. No. M2200L). The ligated DNA was PCR amplified with primers containing Illumina paired-end sequencing adaptors. The final libraries were quantified using bioanalyzer Agilent DNA 1000 (Agilent 5067-1504) and were pooled together in equal molar ratio for paired-end sequencing using MiSeq platform (Illumina) with MiSeq Reagent Kit V3 150-cycle (Illumina).

The sequencing data was de-multiplexed and trimmed to contain only the sgRNA sequence. The sgRNA sequences were mapped to a reference sgRNA library to discard any mismatched sgRNA sequences. The read counts of each sgRNA were calculated. The following analysis was performed with a custom Python script: sgRNAs with read counts less than 50 in the initial time point were discarded; The total read counts were normalized between samples; Artificial one count was assigned to sgRNAs that have zero read count at final time point; The average log2 fold change in abundance of all sgRNA against a given domain/gene was calculated. AML-specific dependency was determined by subtracting the average of log2 fold-change in non-AML cell lines from average log2 fold-change in AML cell lines, and the score was ranked in ascending order. The E2/E3 CRISPR screening data are shown in Supplementary Table 3.

### Analysis of genetic dependencies and gene expression in DepMap and other data sets

Genetic dependency (CRISPR; Avana) data, RNA-seq gene expression (CCLE) data and DNA methylation data (CCLE, promoter 1kb upstream TSS) from cancer cell lines were extracted from the DepMap Public Project Achilles 20Q2 database (http://depmap.org/portal/). RNA expression data with genomic information in AML patient and other tumor patient samples was extracted from The Cancer Genome Atlas database via cBioportal (https://www.cbioportal.org/). Pearson correlation coefficient was calculated in RStudio 1.2.5.

### Competition-based cell proliferation assay

Cas9-expressing cell lines were lentivirally transduced with LRG2.1T sgRNA linked with GFP or mCherry reporter. Percentage of GFP positive cell population was measured at day 3 or day 4 as initial time point using a Guava Easycyte HT instrument (Millipore). GFP% (mCherry%) was then measured every two days (for leukemia cell lines) or every three days (for non-leukemia cell lines) over a time course. The relative change in the GFP% (or mCherry%) percentage at each time point was then normalized to initial time point GFP% (or mCherry%). This relative change was used to assess the impact of individual sgRNAs on cellular proliferation, which reflects cells with a genetic knockout being outcompeted by non-transduced cells in the cell culture.

### Western blot analysis

Cells were collected and washed once with PBS. Cell pellets were resuspended in RIPA buffer (Thermo Scientific; Cat. No. 89901), and sonicated to fragment chromatin. Cell lysate was mixed with SDS-loading buffer containing 2-mercaptoethanol and boiled at 95 °C for 5 min. The cell extracts were separated by SDS-PAGE (NuPAGE 4-12% Bis-Tris protein Gels, Thermofisher), followed by transfer to nitrocellulose membrane using wet transfer at 30 V overnight. Membrane was blocked with 5% nonfat milk in TBST and incubate with primary antibody (1:500 dilution except FLAG antibody which is 1:1000 dilution) in 5% milk in room temperature for one hour. After incubation, membrane was washed for three times with TBST followed by incubation with secondary antibody for one hour at room temperature. After three times wash with TBST, membrane was then incubated with chemiluminescent HRP substrate (Thermo fisher; Cat. No. 34075). Primary antibodies used in this study included UBE2T (abcam; Cat. No. ab140611), FANCD2 (Novus Biotech; Cat. No. NB100-182SS), ALDH1A1 (Proteintech; Cat. No. 22109-1-AP), ALDH2 (Proteintech; Cat. No. 15310-1-AP), FLAG (Sigma; Cat. No. F1804), TP53 (Santa Cruz; sc-126), CDKN1A (Santa Cruz; Cat. No. SC-71811).

### *In vivo* transplantation of MOLM-13 Cells into NSG mice

MOLM-13-Cas9 cells were first transduced with a luciferase expressing cassette in Lenti-luciferase-P2A-Neo (Addgene #105621) vector, followed by G-418 (1 mg/mL) selection and then viral transduction with LRG2.1T-sgRNA-GFP vectors targeting the FANCD2 gene and negative control. Five replicates were performed for each sgRNA. On day 3 post infection with the sgRNA, the infection rate was checked by the percentage of GFP positive cells, and all samples had over 90% infection rate. 0.5 million cells were injected intravenous into sublethally irradiated (2.5Gy) NSG mice (Jax 005557). To detect the disease progression, mice were imaged with IVIS Spectrum system (Caliper Life Sciences) on Day 10, 13, 16 and 19 post injection.

### Cell cycle arrest and apoptosis analysis

Cell cycle analysis was performed according to the manufacture’s protocol (BD, FITC BrdU Flow Kit; Cat. No. 559619), with cells pulsed with BrdU for 1 hour at 37 °C. Cells were co-stained with 4’,6-diamidino-2-phenylindole (DAPI) for DNA content measurement, and analyzed with a BD LSRFortessa flow cytometer (BD Biosciences) and FlowJo software (TreeStar). Annexin V apoptosis staining was performed according to the manufacture’s protocol, with DAPI stained for DNA content measurement (BD, FITC Annexin V Apoptosis Detection Kit; Cat. No. 556547). The experiments were performed in triplicate.

### Immunofluorescence for phospho-H2AX foci

Cells were harvested four days after lentiviral spin-infection, and spun onto a slide using Shandon Cytospin 2 centrifuge. Cells were washed once in PBS, fixed with 3.7% formaldehyde (Sigma) in PBS for 10 min at room temperature, washed twice with PBS, permeabilized with 0.5% Triton X-100 (Sigma) in PBS for 10 min at room temperature, washed in PBS twice, incubated in 5% FBS/PBS for one hour at room temperature for blocking, incubated with primary antibody (anti-phospho-H2AX, clone JBW301; Millipore, Cat. No. 05-636) at 1:1000 dilution at 4 °C overnight, washed with 5% FBS/PBS for 5 min three times, incubated with secondary antibody (anti-moue AF594; Invitrogen, Cat. No. A11005) at 1:2000 dilution for one hour at room temperature, washed with 5% FBS/PBS for 5 min three times, once with PBS and mounted with DAPI Fluoromount-G® (SouthernBiotech). Images were obtained using Zeiss Axio Observer A1 microscope and AxioVision 4.9.1. software. Data were analyzed by CellProfiler 3.1.8.

### RNA-Seq library construction

Total RNA of each cell line was extracted using TRIzol reagent (Thermo Scientific; Cat.No. 15596018) according to the manufacturer’s instructions. Briefly, 3 million cells were lysed with 1 mL of TRIzol and 200 ul chloroform and incubated for 10 min at room temperature followed by centrifuge at 10,000 x g for 15 min at 4 °C. The aqueous phase was added to equal volume of isopropanol and incubated for 10 min at room temperature. RNA was precipitated at 10,000 x g for 10 min at 4 °C, the pellet was washed once with 75% ethanol and dissolved in DEPC-treated water. RNA-Seq libraries were constructed using TruSeq sample preparation kit V2 (Illumina) according to the manufacture’s instruction. Briefly, polyA mRNA was selected from 2 μg of purified total RNA, and fragmented with fragmentation enzyme. First strand of cDNA was synthesized using Super Script II reverse transcriptase, and then second strand was synthesized. Double-stranded cDNA was end-repaired, 3’-adenylated, ligated with indexed adaptor, and then PCR-amplified. The quantity of the RNA-seq library was determined by nanodrop, and the average quantity of RNA-seq libraries ranged from 40 to 80 ng/μL. The same molar amount of RNA-seq library was pooled together and analyzed by single-end sequencing using NextSeq platform (Illumina) with single-end reads of 75 bases.

### RNA-seq data analysis

Sequencing reads were mapped into reference genome hg38 using STAR v2.5.2 with default parameters (Dobin et al., 2013). Read counts tables were created by HTSeq v0.6.1 with customed gtf file containing protein coding genes only. Differentially expressed genes were analyzed using DESeq2 with replicate (Love et al., 2014) using default parameters. RPKMS (reads per kilobase per million mapped reads) were calculated by using Cuffdiff v2.2.1 with default parameters (Trapnell et al., 2013). Genes with RPKMs of more than three in the control were considered as expressed and used in the subsequent analysis. Genes were ranked by their log2 fold change calculated from DESeq2 as input for Pre-ranked GSEA analysis with all available signatures in the Molecular Signature Database v5.2 (MSigDB).

### Aldefluor assay

The Aldefluor assay was performed following the instruction of the Aldefluor Kit (STEMCELL; Cat. No. 01700). Briefly, 0.5 × 10^6^ fresh cell samples were collected and washed once in PBS buffer. The cells were then resuspended in Aldefluor Assay Buffer to 1 × 10^6^/mL. 5 μL of DEAB reagent was added to the cell lines as negative control. 5 μL of activated Aldefluor reagent was added to the control and test samples. The solution was mixed and incubated at 37 °C for 50 min. The cells were collected and resuspended in 500 μL Aldefluor Assay Buffer, and subjected to flow cytometry assay for data acquisition. The experiments were performed in triplicate.

### Mitochondria fractionation

Mitochondrial fractionation was performed using the Mitochondria Isolation Kit for Cultured Cells following the manufacture’s instruction (Thermo Scientific; Cat No. 89874). Briefly, cell membrane was first disrupted by three freeze and thaw cycles, and mitochondria fraction was collected by centrifugation. The collected mitochondrial pellet was resuspended in RIPA buffer for further western blot analysis.

### ChIP-Seq analysis

For ChIP-seq analysis, raw reads were obtained from public GEO datasets MOLM-13(GSE63782), K562 (GSM1652918), MV4-11, THP-1 (GSE79899) and mapped to the human genome (hg19) using Bowtie2 (version 2.2.3) software (Langmead and Salzberg, 2012) using sensitive settings. Duplicate reads were removed prior to peak calling using MACS2 (version 2.1.1.20160309) (Feng et al., 2012) software using 5% FDR cut off and broad peak option. Sequencing depth normalized ChIP-seq pileup tracks were visualization using the UCSC genome browser (Kent et al.).

### Nanopore sequencing

crRNA guides specific to the regions of interest (ROI) were designed as per recommended guidelines described in the Nanopore infosheet on Targeted, amplification-free DNA sequencing using CRISPR/Cas (Version: ECI_S1014_v1_revA_11Dec2018). Guides were reconstituted to 100 μM using TE (pH 7.5) and pooled into an equimolar mix. For each distinct sample, four identical reactions were prepared parallelly using 5 μg gDNA each. Ribonucleoprotein complex (RNPs) assembly, genomic DNA dephosphorylation, and Cas9 cleavage were performed as described in (Gilpatrick et al., 2019). Affinity-based Cas9-Mediated Enrichment (ACME) using Invitrogen™ His-Tag Dynabeads™ was performed to pulldown Cas9 bound non-target DNA, increasing the proportion of on-target reads in the sample (Iyer et al. 2020). The resultant product was cleaned up using 1X Ampure XP beads (Beckman Coulter; Cat. No. A63881), eluted in nuclease-free water, and pooled together. The ACME enriched sample was quantified using Qubit fluorometer (Thermo Fisher Scientific) and carried forward to the adapter ligation step as described by Iyer et al. Sequencing adaptors from the Oxford Nanopore Ligation Sequencing Kit (ONT; SQK-LSK109) were ligated to the target fragments using T4 DNA ligase (NEBNext Quick Ligation Module E6056). The sample was cleaned up using 0.3X Ampure XP beads (Beckman Coulter; Cat. No. A63881), washed with long-fragment buffer (LFB; ONT, SQK-LSK109), and eluted in 15 μL of elution buffer (EB; ONT, LSK109) for 30 min at room temperature. Resultant library was prepared for loading as described in the Cas-mediated PCR-free enrichment protocol from ONT (Version: ENR_9084_v109_revH_04Dec2018) by adding 25 μL sequencing buffer (SQB; ONT, LSK109) and 13 μL loading beads (LB; ONT, LSK109) to 12 μL of the eluate. Each sample was run on a FLO-MIN106 R9.4.1 flow cell using the GridION sequencer.

Real time basecalling was performed with Guppy v3.2, and files were synced to our Isilon 400NL storage server for further processing on the shared CSHL HPCC. Nanopolish v0.13.2 (Simpson et al., 2017) was used to call methylation per the recommended workflow. Briefly, indexing was performed to match the ONT fastq read IDs with the raw signal level fast5 data. The ONT reads were then aligned to the human reference genome (UCSC hg38) using minimap2 v2.17 (Li, 2018) and the resulting alignments were then sorted with samtools v0.1.19 (Li et al., 2009). Nanopolish call-methylation was then used to detect methylated bases within the targeted regions – specifically 5-methylcytosine in a CpG context. The initial output file contained the position of the CpG dinucleotide in the reference genome and the methylation call in each read. A positive value for log_lik_ratio was used to indicate support for methylation, using a cutoff value of 2.0. The helper script calculate_methylation.py was then used to calculate the frequency of methylation calls by genomic position.

### RT-qPCR analysis following 5-azacytidine treatment

For dose-dependent experiment, 5-azacytidine was added to cell culture with different concentrations, and cells were collected after 36 h treatment. For time-course treatment, cells were treated with 1 μM 5-azacytidine and collected at different time points. Total RNA was extracted using TRIzol reagent as described above. 1-2 μg of total RNA was treated with DNaseI and reverse transcribed to cDNA using qScript cDNA SuperMix (Quantabio; Cat. No. 84033), followed by RT-qPCR analysis with SYBR green PCR master mix (Thermo Fisher; Cat. No. 4309155) on a QuantStudio™ 7 Flex Real-Time PCR System. GAPDH was used as reference gene. The primers used in this study are listed in Supplementary Table 2.

### Aligned enhanced reduced representation bisulfite sequencing (ERRBS)

myCpG files from AML (n=119) and normal (n=22) individuals were downloaded from GEO (accession number GSE98350) (Glass et al., 2017). After filtering and normalizing by coverage, a methylBase object containing the methylation information and locations of cytosines that were present in at least 5 samples per condition (meth.min=5) was generated using MethylKit (version 1.9.3) (Akalin et al., 2012) and R statistical software (version 3.5.1). Percent methylation for each CG for each donor, was calculated using the MethylKit ‘percMethylation’ function. Bedtools (Quinlan and Hall, 2010) intersect function was used to determine overlap with CpGi from hg19. The heatmap of the percent methylation of the cytosines covered within the ALDH2 CpGi was plotted using ComplexHeatmap (Gu et al., 2016), with the complex clustering method and Euclidian distances, and using light grey for NA values. For the boxplot of the average percent methylation for each sample at the CpGi, the mean methylation for each sample was calculated using all CG that were covered in that region, using R. Data was plotted using ggplot2 and significance calculated using the ggplot2 function ‘stat_compare_means’ with the Student’s t-test method. To correlate methylation with gene expression, processed expression data that was generated for the above samples using the Affymetrix Human Genome 133 Plus2.0 GeneChips was downloaded (Glass et al., 2017; Verhaak et al., 2009). The average percent methylation of the regions covered within the CpGi versus RNA expression for ALDH2 was plotted using ggplot2 and fitted with a line calculated with a linear model. Pearson’s correlation method was used to determine significance.

### Statistics

Error bars represent the mean plus or minus standard error of the mean, and n refers to the number of biological repeats. Statistical significance was evaluated by p value using GraphPad Prism software as indicated in the figure legends. For Kaplan-Meier survival curves, the log rank (Mantel-Cox) test was used to estimate median overall survival and statistical significance.

## RESULTS

### A domain-focused CRISPR screen targeting the ubiquitination machinery identifies UBE2T and FANCL as unique AML dependencies

In this study, we pursued the identification of AML-specific dependencies on the ubiquitination machinery using domain-focused CRISPR sgRNA screening, which is a strategy for profiling the essentiality of protein domain families in cancer cell lines (Shi et al., 2015). Using an sgRNA design algorithm linked to protein domain annotation, we designed 6,079 sgRNAs targeting exons encoding 573 domains known to be involved in ubiquitin conjugation or ligation, which were cloned in a pooled manner into the LRG2.1T lentiviral vector (Figure 1A; Table S1). Using this sgRNA library, we performed negative selection “dropout” screening in twelve Cas9-expressing human cancer cell lines, which included six AML and six solid tumor cell lines (Figure 1B). Many of the dependencies identified in these screens were pan-essential across the twelve lines, such as ANAPC11, CDC16, and RBX1 (Figure S1A). We ranked all ubiquitination-related genes based on their degree of essentiality in AML versus solid tumor contexts, which nominated UBE2T and FANCL as AML-specific dependencies (Figure 1C and D). The known function of UBE2T and FANCL as E2 and E3 proteins, respectively, within the Fanconi anemia (FA) pathway suggested a unique necessity of this DNA repair function in AML (Ceccaldi et al., 2016; Kottemann and Smogorzewska, 2013).

**Figure 1.**
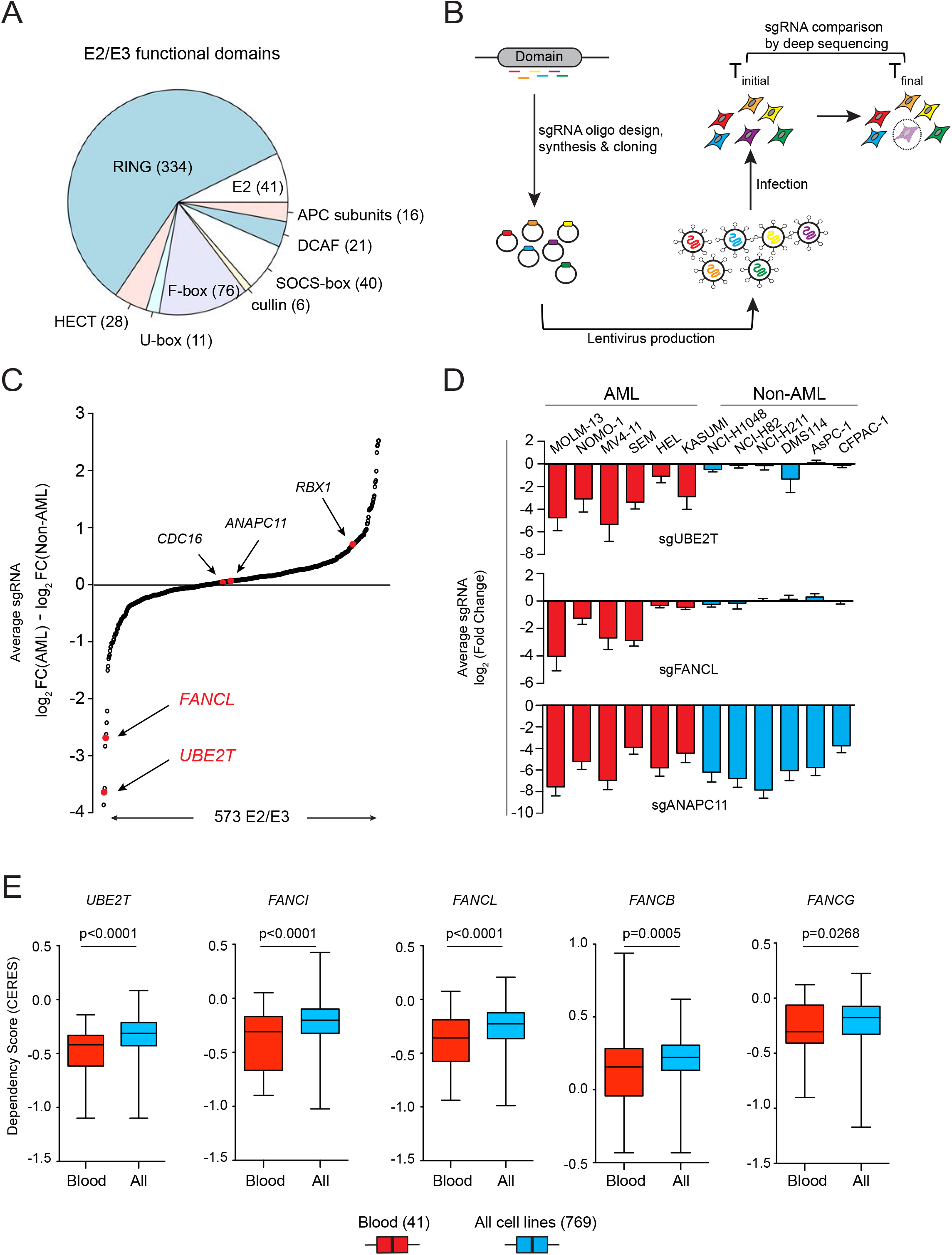
Domain-Focused CRIPSR screening identifies UBE2T and FANCL as AML-biased dependencies. **(A)** Categories of E2 and E3 domains targeted in the ubiquitination domain sgRNA library. sgRNA sequences are provided in Supplementary Table 1. **(B)** Schematic of CRISPR-Cas9 screening strategy. **(C)** Ubiquitination machinery dependencies ranked by the average log2 fold-change difference comparing six AML and six non-AML cell lines in the pooled CRISPR screening. **(D)** Average log2 fold-change for sgRNAs targeting UBE2T, FANCL, and ANAPC11 in the twelve screened cell lines. **(E)** Dependency scores of five FA genes extracted from Project Achilles (20Q2) (Dempster et al., 2019; Meyers et al., 2017). The boxplots indicate the distribution of dependency score (CERES; a normalized metric of gene essentiality) of five FA genes across all 769 cell lines or in the subset of 41 blood cell lines. p-value was calculated using an unpaired Student’s t-test. All sgRNA experiments were performed in Cas9-expressing cell lines.

We corroborated the biased essentiality of UBE2T/FANCL in blood cancers relative to other cancer types by analyzing data obtained from Project Achilles (version 20Q2), in which genome-wide CRISPR essentiality screening was performed in 729 cancer cell lines (Figure 1E; Figure S1B) (Dempster et al., 2019; Meyers et al., 2017). In addition, these data revealed that several other FA pathway genes, including FANCI, FANCB and FANCG, are also blood cancer-biased dependencies in a manner that correlated with UBE2T/FANCL essentiality. These findings reinforce that blood cancer cells are hypersensitive to perturbation of FA proteins relative to other cancer types.

### FA proteins are dependencies in a subset of AML cell lines under *in vitro* and *in vivo* conditions

To further validate the specificity of FA protein dependencies in AML, we performed sgRNA competition assays following inactivation of UBE2T, FANCL, and FANCD2 genes in 27 human cancer cell lines, including 14 leukemia, 5 pancreatic cancer, 4 lung cancer, and 4 sarcoma lines (Figure 2A and B; Figure S2A and B). These experiments showed that targeting any of these three FA genes suppressed the fitness of 9 human leukemia lines, including 3 generated by retroviral transduction of oncogenes into human hematopoietic stem and progenitor cells (Wei et al., 2008). In contrast, the fitness of 5 other human leukemia lines and all of the solid tumor cell lines was less sensitive to targeting of FA genes (Figure 2A). As a positive control for this assay, targeting of CDK1 arrested the growth of all cancer cell lines tested. Western blotting confirmed that the variable pattern of growth arrest following UBE2T targeting was not due to differences in genome editing efficiency (Figure 2B; Figure S2A).

**Figure 2.**
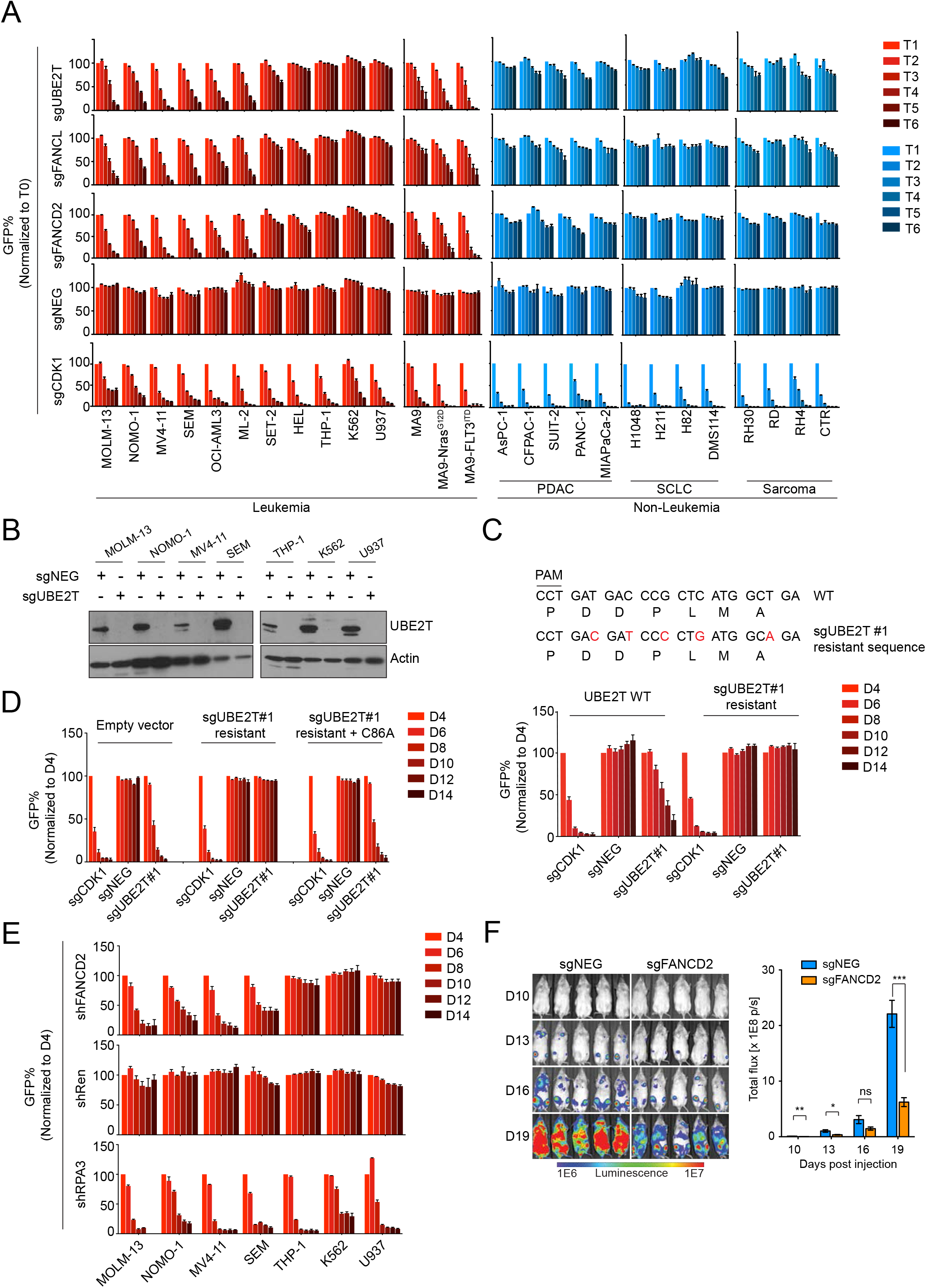
FA proteins are AML-biased dependencies *in vitro* and *in vivo*. **(A)** Competition-based proliferation assays performed in 27 human cancer cell lines by lentiviral infection with the indicated sgRNAs linked to GFP. Timepoints (T) were collected every 2 days for leukemia cells or every 3 days for non-leukemia lines. Rate of GFP depletion indicate loss of cell fitness caused by Cas9/sgRNA-mediated genetic mutations. PDAC: Pancreatic Ductal Adenocarcinoma; SCLC: Small Cell Lung Cancer. (n=3). **(B)** Western blotting performed on whole cell lysates prepared from indicated cell lines, performed on day 4 following lentiviral transduction with indicated sgRNAs. **(C)** Competition-based proliferation assays comparing impact of sgRNAs (linked with GFP) on MOLM-13 cell fitness, in the context of co-transduction with UBE2T WT or sgRNA-resistant UBE2T cDNA. The sgRNA-resistant cDNA mutagenesis strategy is depicted. (n=3). **(D)** Competition-based assays measuring sgRNA (linked to GFP) effects on MOLM-13 cell fitness, in the presence of the indicated cDNAs. (n=3). **(E)** Competition-based proliferation assays in MOLM-13 cells lentivirally infected with indicated shRNA (linked to GFP) targeting FANCD2. (n=3). **(F)**(Left) Bioluminescence imaging of NSG mice transplanted with luciferase+/Cas9+ MOLM-13 cells infected with either sgNEG or sgFANCD2. (Right) Quantification of bioluminescence intensity. (n=5). All bar graphs represent the mean ± SEM. p-value is calculated by unpaired Student’s t-test. ***p < 0.001, **p < 0.01, *p < 0.05. All sgRNA experiments were performed in Cas9-expressing cell lines.

As additional controls, we verified that the growth arrest caused by UBE2T or FANCL knockout in MOLM-13 cells was due to an on-target effect by rescuing this effect with a cDNA carrying silent mutations that abolish sgRNA recognition (Figure 2C; Figure S2C-F). Using this rescue assay, we investigated whether the catalytic function of UBE2T was essential for AML growth by comparing the wild-type cDNA with the C86A mutation, which abolishes ubiquitin conjugation activity (Alpi et al., 2008; Machida et al., 2006). Despite being expressed at a similar level to the wild-type protein, the C86A mutant of UBE2T was unable to support AML growth, suggesting that the ubiquitination cascade involving UBE2T-FANCL is essential in this context (Figure 2D, Figure S2G and H). To ensure that this dependency was not influenced by DNA damage caused by CRISPR-Cas9, we also validated FANCD2 dependency in AML using RNAi-based knockdown (Figure 2E; Figure S2I). In addition, inactivation of FANCD2 suppressed the growth of MOLM-13 cells when propagated *in vivo* in immune deficient mice (Figure 2F; Figure S2J).

### Targeting of FA proteins in AML leads to cell cycle arrest and apoptosis in a p53-dependent manner

We next performed a deeper characterization of the AML cell phenotype following inactivation of FA proteins. A flow cytometry analysis of BrdU incorporation and DNA content revealed an accumulation of UBE2T- and FANCD2-deficient MOLM-13 cells in G1 and in G2/M phase, with a significant loss of cells in S phase (Figure 3A and B). An Annexin V staining analysis of UBE2T- or FANCD2-deficient MOLM-13 cells also revealed evidence of apoptosis (Figure 3C and D). Levels of γH2AX, a marker of DNA damage, were also increased following FA gene inactivation (Figure S3A). As apoptosis and cell cycle arrest are known downstream consequences of DNA damage-induced p53 activation, we investigated this pathway in FA-deficient AML cells using RNA-seq in MOLM-13 cells. Inactivation of UBE2T, FANCL or FANCD2 led to significant upregulation of p53 target genes (Figure 3E; Figure S3B). In addition, we observed induction of pro-apoptotic genes and suppression of DNA replication genes following FA protein inactivation, in accord with the phenotypes described above (Figure 3E; Figure S3B). To evaluate the contribution of p53 to FA dependence in AML, we inactivated *TP53* or its target gene *CDKN1A* using CRISPR genome editing in MOLM-13 or MV4-11, which rendered cells less sensitive to loss of UBE2T (Figure 3F and G; Figure S3C-F). These findings indicate that loss of FA proteins in AML triggers cell cycle arrest and apoptosis in a p53-dependent manner.

**Figure 3.**
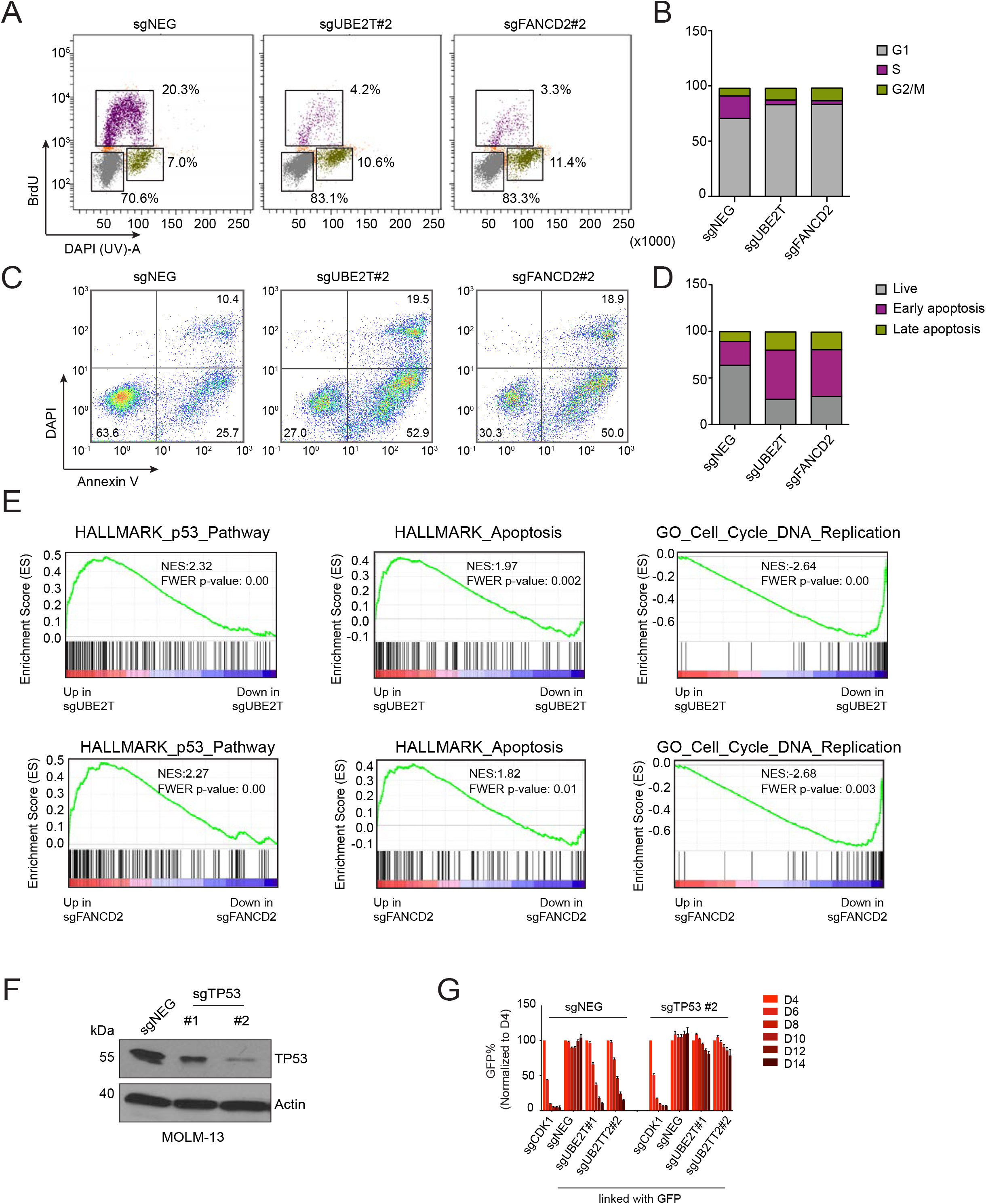
Inactivation of Fanconi anemia genes in AML leads to p53-induced cell cycle arrest and apoptosis. **(A)** Representative flow cytometry analysis of BrdU incorporation and DNA content to infer cell status following lentiviral transduction of MOLM-13 cells with the indicated sgRNAs (day 6). **(B)** Quantification of different cell cycle stages shown in (A). **(C)** Representative flow cytometry analysis of DAPI (indicating permeable dead cells) and annexin-V staining (a pre-apoptotic cell marker) following lentiviral transduction of MOLM-13 cells (day 6). **(D)** Quantification of live and apoptotic cells shown in (C). **(E)** Gene set enrichment analysis (GSEA) of RNA-seq data obtained from MOLM-13 cells lentivirally transduced with the indicated sgRNAs (Subramanian et al., 2005). Normalized enrichment score (NES) and family-wise error rate (FWER) *p*-value are shown. **(F)** Western blot analysis performed on lysates obtained from MOLM-13 cells on day 6 following sgRNA transduction. **(G)** Competition-based proliferation assays in MOLM-13 cells following sequential sgRNA transduction. Negative sgRNA or TP53 sgRNAs were infected first, selected with neomycin, followed by transduction with the sgRNAs indicated at the bottom of the graph (linked with GFP). n=3. All bar graphs represent the mean ± SEM. All sgRNA experiments were performed in Cas9-expressing cell lines.

### Low Aldefluor activity and ALDH1A1/ALDH2 expression in FA-dependent AML cell lines

We next hypothesized that an endogenous source of DNA damage might drive the elevated demand for FA proteins in AML. For example, excess accumulation of aldehydes can exacerbate the phenotypes of FA-deficient mice and humans (Garaycoechea et al., 2012; Hira et al., 2013; Langevin et al., 2011). This led us to investigate the status of aldehyde dehydrogenases (ALDH), which are comprised of 19 enzymes that oxidize diverse aldehydes into non-toxic acetates in a NAD- or NADP-dependent manner (Langevin et al., 2011). We first made use of the Aldefluor assay, in which cells are treated with BODIPY-aminoacetaldehyde (BAAA), a cell-permeable fluorescent aldehyde that becomes trapped in cells following ALDH-mediated conversion to BODIPY-aminoacetate (BAA). After applying BAAA to a diverse collection of human AML cell lines, we observed that most FA-dependent lines exhibited minimal ALDH activity, with fluorescence levels similar to control cells treated with the pan-ALDH inhibitor N,N-diethylaminobenzaldehyde (DEAB). In contrast, ALDH activity was higher in most of the FA-dispensable group of AML lines (Figure 4A). One exception to this correlation was U937 cells, which are FA-dispensable but lacks ALDH activity. However, we note that U937 lacks a functional p53 pathway (Ghandi et al., 2019), which would be expected to alleviate the FA-dependence in this context.

**Figure 4.**
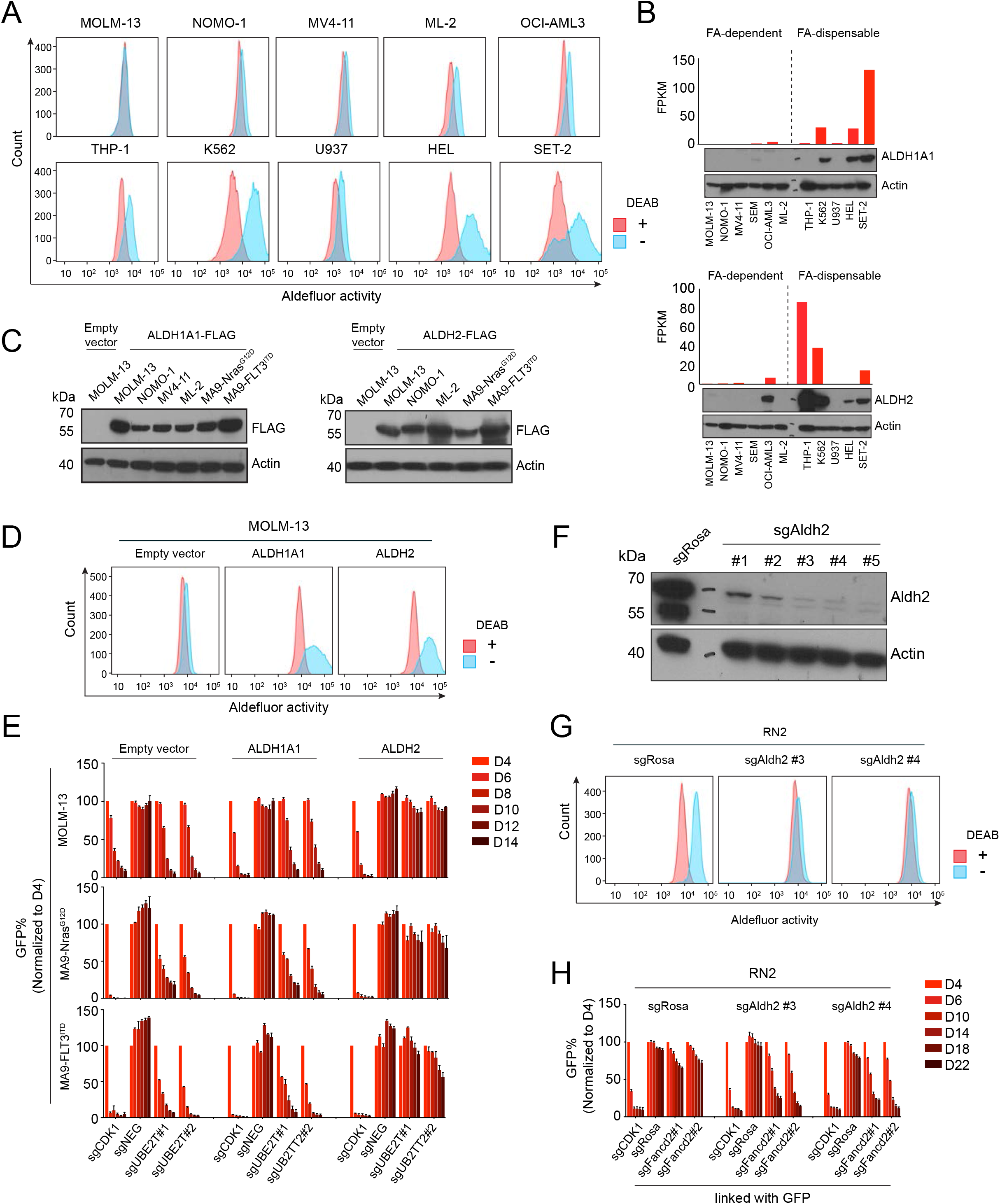
Silencing of ALDH2 expression in AML confers Fanconi anemia proteins dependency. **(A)** Representative flow cytometry analysis of Aldefluor assays, in which AML cell lines were treated with 3.75 μM BAAA, with or without the ALDH inhibitor DEAB (15 μM). **(B)** RNA and protein levels of *ALDH1A1* or *ALDH2* measured by RNA-seq (top) and western blot (bottom) in the indicated cell lines. **(C)** Western blot analysis of whole cell lysates from cell lines transduced with the indicated lentiviral cDNA constructs. **(D)** Representative Aldefluor analysis of MOLM-13 cells expressing FLAG-tagged ALDH1A1 or ALDH2. **(E)** Competition-based proliferation assay in MOLM-13 expressing the indicated cDNAs, followed by transduction with GFP-linked sgRNA. **(F)** Western blot analysis of ALDH2 expression in RN2 cells following lentiviral transduction with the indicated sgRNAs. **(G)** Aldefluor assay performed in RN2 cells following lentiviral transduction with the indicated sgRNAs. **(H)** Competition-based proliferation assays performed in RN2 cells (control vs Aldh2 knockout) with the indicated sgRNAs linked to GFP. All bar graphs represent the mean ± SEM, (n=3). All sgRNA experiments were performed in Cas9-expressing cell lines.

An RNA-seq analysis revealed that among the 19 ALDH enzymes, only *ALDH1A1* and *ALDH2* were expressed at reduced levels in the FA-dependent lines when compared to FA-dispensable lines (Figure 4B; Figure S4A). Using western blotting, we confirmed that ALDH1A1 and ALDH2 proteins are present at higher levels in the FA-dispensable versus the FA-dependent AML lines (Figure 4B). Of note, while ALDH1A1 and ALDH2 are both capable of oxidizing BAAA, each enzyme is known to have distinct cellular functions (Chen et al., 2014, 2012; Fan et al., 2003; Fernandez et al., 2006; Kitagawa et al., 2000; Ohsawa et al., 2008; Tomita et al., 2016). ALDH1A localizes in the cytosol and is known to oxidize retinol aldehydes into retinoic acids (Fan et al., 2003). In contrast, ALDH2 is localized in the mitochondria and oxidizes acetaldehyde (derived from exogenous ethanol) and 4-Hydroxynonenal (4-HNE), an endogenous aldehyde derived from lipid peroxidation (Chen et al., 2014).

### Re-expression of ALDH2, but not ALDH1A1, renders FA proteins dispensable for AML growth

Considering the inverse correlation between ALDH1A1/ALDH2 expression and FA-dependence in AML lines, we next performed experiments to explore a synthetic lethal genetic interaction in this context. We lentivirally transduced the FA-dependent MOLM-13 line with ALDH1A1 or ALDH2 cDNA and confirmed protein expression via western blotting (Figure 4C). Aldefluor assays revealed that the lentivirally expressed proteins were enzymatically active, with both ALDH1A1 and ALDH2 causing oxidation of BAAA (Figure 4D), consistent with prior findings (Garaycoechea et al., 2012; Nakahata et al., 2015). We next used CRISPR to target UBE2T in the ALDH1A1- or ALDH2-expressing MOLM-13 cells, followed by competition-based assays to track changes in cell fitness. Remarkably, the ALDH2-expressing MOLM-13 cells became resistant to growth arrest caused by UBE2T inactivation, whereas ALDH1A1-expressing cells remained sensitive to UBE2T targeting (Figure 4E). To evaluate the generality of this result, we expressed ALDH1A1 and ALDH2 in four other FA-dependent AML contexts, which likewise showed that ALDH2, but not ALDH1A1, alleviated the dependency on FA proteins (Figure 4E; Figure S4B and C). To address whether inactivation of ALDH2 is sufficient to confer FA dependence in AML, we made use of the murine AML cell line RN2, which was derived by retroviral transformation of hematopoietic stem and progenitors cells with the MLL-AF9 and Nras^G12D^ cDNAs (Zuber et al., 2011). Notably, this cell line retains an intact p53 pathway and expresses ALDH2, but not ALDH1A1 (Figure S4D). We targeted ALDH2 using CRISPR in this cell line and confirmed loss of protein expression and loss of Aldefluor activity (Figure 4F and G). Notably, in competition-based cell fitness assays we observed that ALDH2-deficient RN2 cells became hypersensitive to the inactivation of FANCD2 relative to the parental cells (Figure 4H). These experiments suggest that loss of ALDH2 confers dependency on FA proteins in AML.

We next investigated whether the catalytic function of ALDH2 was needed for the bypass of FA pathway dependency by comparing the wild-type ALDH2 cDNA with the E268K and the C302A alleles (Nene et al., 2017). Importantly, these two mutant proteins were expressed normally, but lacked any detectable Aldefluor activity. In addition, both mutants were unable to rescue the UBE2T dependency of MOLM-13 cells (Figure S4E-H). As an additional control, we considered whether ALDH1A1 might be capable of rescuing the FA dependency if we forced its localization into the mitochondria using two different localization signals (Figure S5A). Despite effective targeting to the mitochondria confirmed by cell fractionation and robust Aldefluor activity of these mutant proteins (Figure S5B and C), the mitochondrial forms of ALDH1A1 were unable to rescue UBE2T dependence (Figure S5D). Other ALDH enzymes (ALDH1A2, ALDH1B1, ALDH6A1, ALDH7A, or ALDH1A3) were also tested in this assay, but were unable to rescue the UBE2T dependence when lentivirally expressed in MOLM-13 cells (Figure S5E-H). Taken together, these experiments suggest a unique capability of ALDH2 to detoxify a specific subset of endogenous aldehydes that drive FA protein dependency in AML.

Interestingly, restoration of ALDH2 expression in MOLM-13 cells led to no detectable impact on cell proliferation *in vitro* (Figure S6A). In addition, an RNA-seq analysis suggested that the overall transcriptome of MOLM-13 cells was largely unaffected by re-expression of ALDH2 (Figure S6B). These findings suggest that loss of ALDH2 in AML leads to a discrete defect in aldehyde detoxification, which is largely compensated for by the presence of the FA DNA repair pathway.

### Silencing and hypermethylation of *ALDH2* occurs in a recurrent manner in human AML

We next investigated whether *ALDH2* silencing is associated with DNA hypermethylation, which is known to be aberrantly distributed across the AML genome (Glass et al., 2017). In AML cell lines profiled in the Cancer Cell Line Encyclopedia Project (Barretina et al., 2012; Ghandi et al., 2019), we detected an inverse correlation between levels of DNA methylation and *ALDH2* expression (Figure 5A). To confirm this finding, we analyzed DNA methylation at the *ALDH2* promoter in MOLM-13 and MV4-11 cells using Nanopore sequencing (Simpson et al., 2017), which confirmed dense hypermethylation in the vicinity of the *ALDH2* promoter (Figure 5B). Importantly, this same location was hypomethylated in normal human hematopoietic stem and progenitor cells (Figure 5B) (Hodges et al., 2011). In addition, silencing of *ALDH2* correlated with diminished histone acetylation at this genomic region (Figure 5C). Using an inhibitor of DNA methyltransferase activity 5-azacytidine, we confirmed a time- and dose-dependent increase in *ALDH2* expression in MOLM-13 and MV4-11 cells following compound exposure (Figure 5D and E; Figure S7A and B). These results suggest that silencing of *ALDH2* in AML cell lines is associated with the acquisition of DNA hypermethylation.

**Figure 5.**
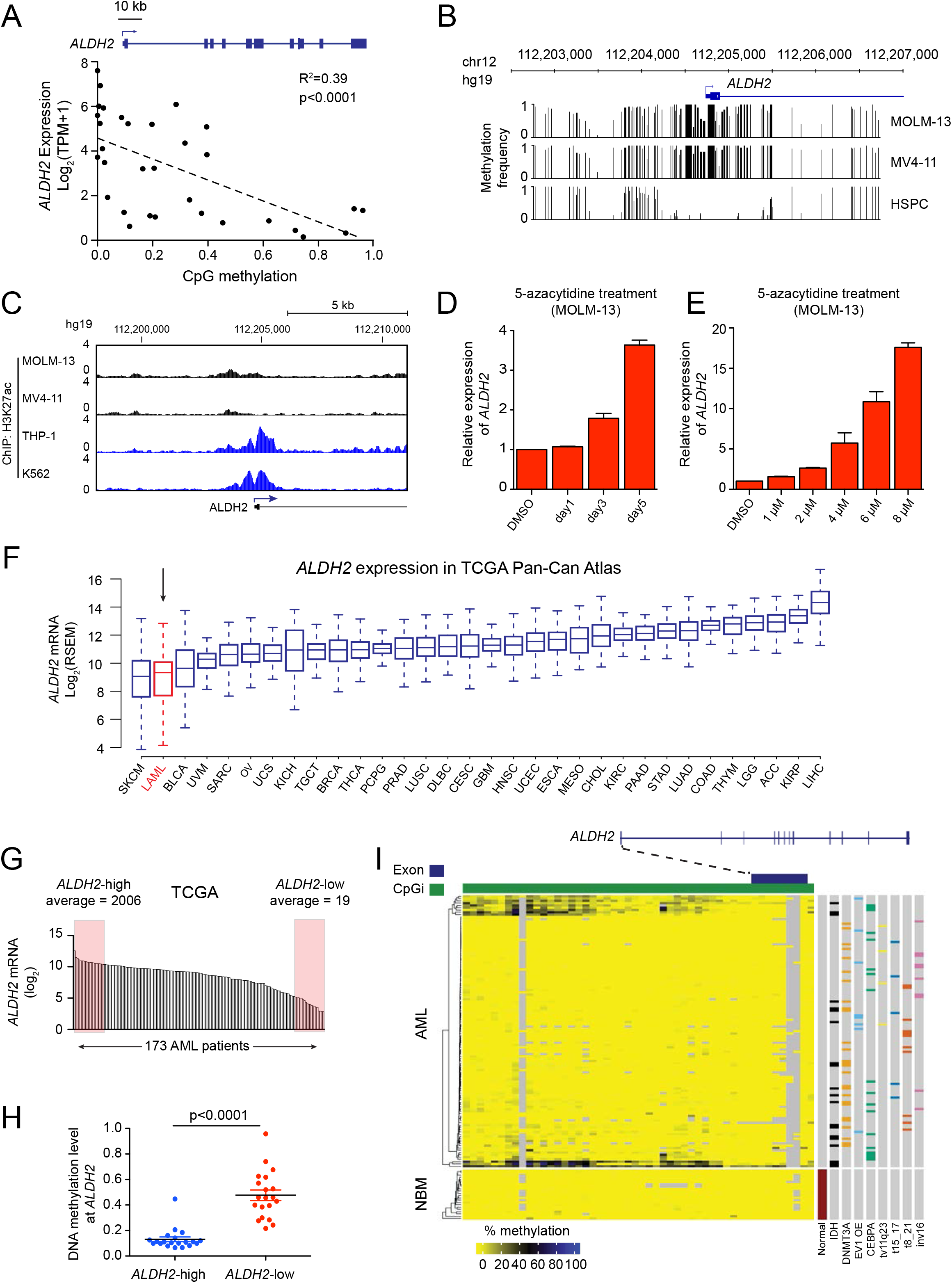
Silencing of *ALDH2* in human AML cell lines and patient samples is associated with promoter DNA hypermethylation. **(A)** Scatterplot analysis comparing *ALDH2* mRNA levels and DNA methylation levels at the *ALDH2* promoter across 32 AML cell lines (data was obtained from the Cancer Cell Line Encyclopedia) (Barretina et al., 2012; Ghandi et al., 2019). p value is calculated by linear regression. **(B)** DNA methylation analysis of the *ALDH2* promoter in leukemia cell lines (Nanopore sequencing) or in normal human hematopoietic stem and progenitor cells (bisulfite sequencing from (Hodges et al., 2011)). **(C)** ChIP-seq analysis of H2K27ac enrichment at the *ALDH2* promoter. **(D-E)** RT-qPCR analysis of ALDH2 mRNA levels in MOLM-13 cells following treatment with 5-azacytidine. In (D), 1 μM concentration was used. In (E), a 36 hour timepoint was used. **(F)** ALDH2 mRNA levels extracted from the TCGA Pan-Can Atlas (The Cancer Genome Atlas Research Network, 2013). **(G)** ALDH2 mRNA levels in 173 AM patient samples from TCGA. Shown are the samples classified as ALDH2-high and ALDH2-low. **(H)** Comparison of CpG DNA methylation level at the ALDH2 promoter of indicated AML patient samples. p-value is calculated by unpaired Student’s t-test. **(I)** Heatmap of the percent methylation of the covered CG within the *ALDH2* promoter. Each column is one CG and each row is one patient. Rows are clustered using Euclidian distances and the complex clustering method and missing CGs are depicted in gray. Select patient mutational and cytogenic data is plotted on the right of the heatmap. All bar graphs represent the mean ± SEM (n=3).

We next investigated whether silencing and hypermethylation of *ALDH2* occurs in human AML patient specimens. By analyzing RNA-seq data obtained from diverse tumors included in The Cancer Genome Atlas (TCGA) pan-cancer analysis, we found that AML had greater variability in *ALDH2* expression when compared to most other human cancer types (Figure 5F). Within this group of TCGA patient samples, we designated ALDH2-low and ALDH2-high AML, distinguished by a ~100-fold difference in ALDH2 expression (Figure 5G). Notably, we observe that a housekeeping gene *ACTB* only shows a 4-fold variance in expression across TCGA AML samples (Figure S7C). In accord with cell line observations, ALDH2-low AML samples in the TCGA possessed elevated levels of DNA methylation at *ALDH2* relative to ALDH2-high AML samples (Figure 5H). We further confirmed the aberrant hypermethylation and silencing of *ALDH2* in a subset of AML in an independent collection of patient samples (Glass et al., 2017), whereas normal bone marrow cells remained hypomethylated (Figure 5I; Figure S7D-E). Together, these findings suggest that DNA hypermethylation and silencing of *ALDH2* occurs in a recurrent manner in human AML.

## DISCUSSION

Using a genetic screen focused on the ubiquitination machinery, we uncovered a role for FA proteins as dependencies in AML. We account for this observation by the aberrant expression of ALDH2, an enzyme that oxidizes aldehydes in normal tissues but becomes epigenetically silenced in this disease context. We propose that silencing of *ALDH2* in AML leads to an accumulation of endogenous aldehydes, which in turn leads to the formation of DNA crosslinks that necessitate repair by FA proteins. Upon inactivation of FA genes in ALDH2-deficient AML, the levels of aldehyde-induced DNA damage reach a threshold that triggers p53-mediated cell cycle arrest and programmed cell death. This study reinforces how aberrant gene silencing can disable redundant pathways and can lead to acquired dependencies in cancer.

The synthetic lethal interaction between ALDH2 and FA genes is well-supported by observations in mice, in which the combined deficiency of *Aldh2* and *Fancd2* leads to developmental defects, a predisposition to cancer, and a hypersensitivity to exogenous aldehydes (Langevin et al., 2011). These phenotypes were thought to be uniquely present in normal hematopoietic stem and progenitor cells (Garaycoechea et al., 2012), but our study now shows that this genetic interaction extends to malignant myeloid cells. In normal mouse hematopoietic stem cells, aldehyde-induced DNA damage leads to the formation of double-stranded DNA breaks, which ultimately causes stem cell attrition, a phenotype that resembles the clinical presentation of germline FA deficiency in humans (Garaycoechea et al., 2018). Similar to our observations in AML, hematopoietic stem cells in *Aldh2/Fancd2* compound deficient mice become cleared through a p53-dependent mechanism (Garaycoechea et al., 2018). The redundant function of ALDH2 and FA proteins is also supported by evidence in humans, in which a combined hereditary deficiency of *ALDH2* and FA genes correlates with an accelerated rate of bone marrow failure when compared to FA patients with intact ALDH2 function (Hira et al., 2013). Thus, our study builds upon prior work by revealing a clinical context in which the ALDH2/FA protein redundancy is disabled in humans and could be exploited to eliminate AML *in vivo*.

Several prior studies have applied Aldefluor assays to human clinical samples and observed diminished ALDH activity in AML when compared to normal hematopoietic stem and progenitor cells, in accord with our own findings (Gasparetto et al., 2017; Gerber et al., 2012; Hoang et al., 2015; Schuurhuis et al., 2013). Loss of ALDH1A1 expression was previously observed in AML, which correlated with favorable prognosis and was found to render cells hypersensitive to toxic ALDH substrates, such as arsenic trioxide (Gasparetto et al., 2017). Our study validates ALDH1A1 silencing as a recurrent event in AML cell lines, however our functional experiments demonstrate that this event is unrelated to FA dependency in this disease. Collectively, our study and the work of Gasparetto et al point to distinct functional consequences upon loss of ALDH1A1 versus ALDH2 in AML.

ALDH1A1 and ALDH2 are known to have overlapping aldehyde substrates, however our study points to the existence of endogenous genotoxic aldehydes that are uniquely oxidized by ALDH2 in AML. While the high reactivity of aldehydes precludes us from performing an unbiased assessment of aldehyde species in AML cells, it is known that 4-HNE levels are elevated in Aldh2-deficient mice and humans (Guo et al., 2013; Ohsawa et al., 2008). 4-HNE is a byproduct of lipid peroxidation, and can form toxic adducts with DNA and proteins in a variety of cellular pathologies. (Czerwińska et al., 2014; Voulgaridou et al., 2011). Nevertheless, a key unanswered question in the field is the identity of the endogenous sources of genotoxic aldehydes and the therapeutic utility of their pharmacological modulation in disease.

Our experiments suggest that ALDH2 silencing has no measurable impact on AML cell fitness, owing to compensation via FA proteins. Why then is *ALDH2* recurrently silenced in AML? We propose at least three possibilities. First, loss of ALDH2 expression might be under positive selection to confer a critical metabolic adaptation for the early *in vivo* expansion of an AML clone, while at later stages of AML progression (reflected by our cell lines) ALDH2 silencing is no longer relevant for cell proliferation. A second possibility is that loss of ALDH2 promotes the genetic evolution of AML by increasing the probability of acquiring aldehyde-induced genetic mutations. A third hypothesis is that *ALDH2* is merely a gene susceptible to DNA hypermethylation, with epigenetic silencing occurring as a passenger event in this disease. Irrespective of these different scenarios, our study demonstrates how epigenetic silencing can lead to acquired dependencies in cancer.

Our study reveals a paradox: a germline deficiency of FA genes leads to an elevated risk of AML formation while sporadic AML can acquire a dependency on FA proteins to sustain cell proliferation and survival. The contextual nature of this pathway is reminiscent of other DNA repair regulators, such as ATM, which act to protect normal tissues from cancer-causing somatic mutations while inhibition of this pathway can hypersensitize transformed cells to DNA damaging agents (Cremona and Behrens, 2014; Helleday et al., 2008; Sullivan et al., 2012). Considering the age-dependent onset of symptoms in FA patients, a possibility exists that acute and reversible inhibition of the FA pathway may have a therapeutic index in AML, as has been demonstrated for targeting of other DNA repair proteins (Santos et al., 2014). Therefore, our study provides justification for evaluating pharmacological inhibition of UBE2T/FANCL-mediated ubiquitination as therapeutic approach for eliminating ALDH2-deficient AML.

## ACKNOWLEDGMENTS

We thank James C. Mulloy for sharing genetically engineered human AML cell lines. This work was supported by Cold Spring Harbor Laboratory NCI Cancer Center Support grant 5P30CA045508. Additional funding was provided to C.R.V. by the Pershing Square Sohn Cancer Research Alliance, National Institutes of Health grants R01 CA174793 and P01 CA013106, a Leukemia & Lymphoma Society Scholar Award. A. S. is an HHMI Faculty Scholar and is supported by CA204127 from NIH and K99 HL150628 (M.J).

## AUTHORSHIP CONTRIBUTIONS

Z.Y., Y.W., X.S.W, S.V.I., M.K., O.K., O.E.D., K.C., M.E.F., S.G., performed experiments and/or analyzed the data; M.E.F., W.R.M, E.H., A.S., and C.R.V. supervised the experiments and analysis; Z.Y. and C.R.V. wrote the manuscript.

## SUPPLEMENTAL FIGURE LEGENDS

**Figure S1.**
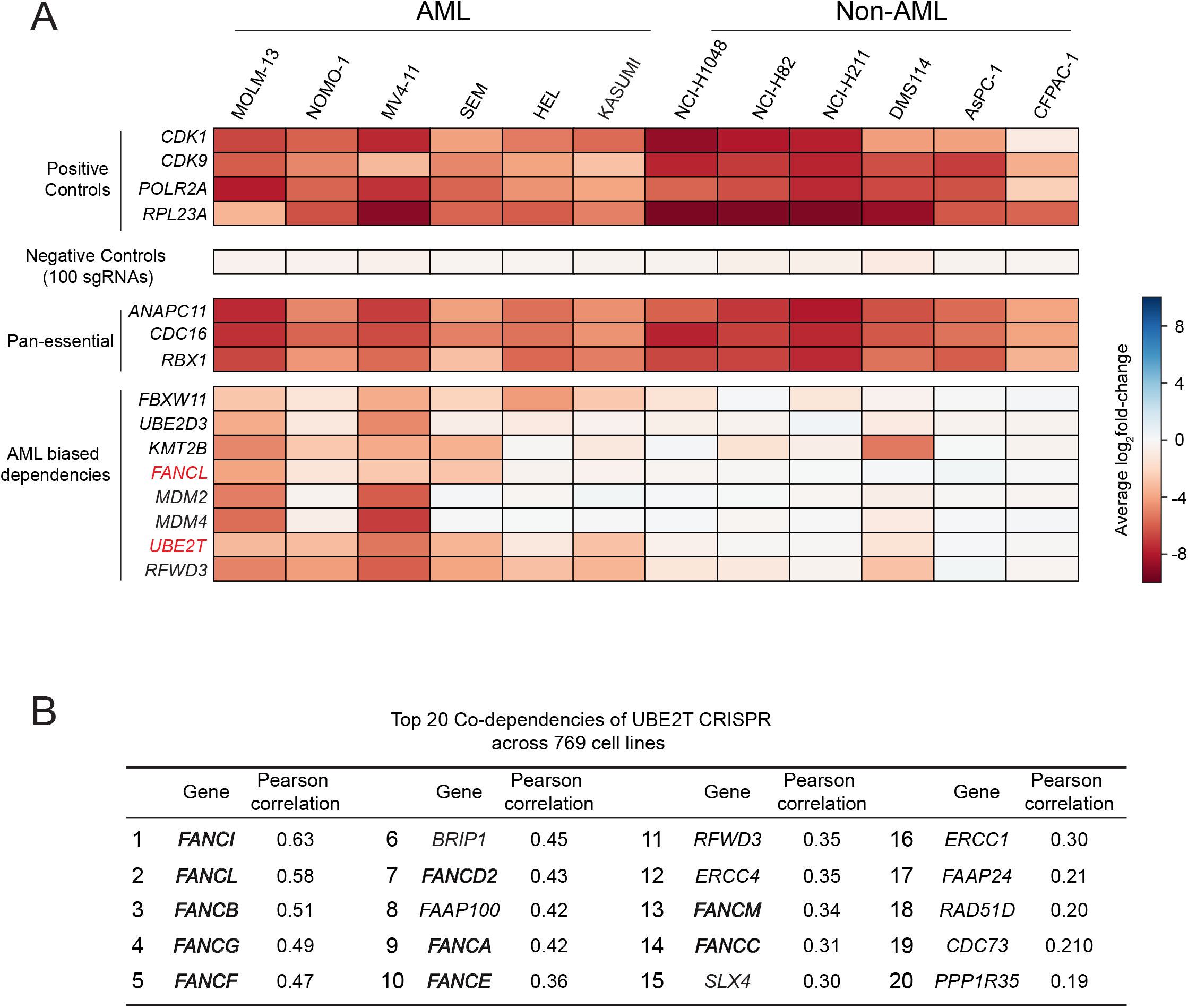
Domain-focused CRISPR screening identifies UBE2T and FANCL as AML dependencies. Related to Figure 1. **(A)** Summary of the fold-depletion of spike-in positive and negative control sgRNAs in the pooled ubiquitination domain-focused CRISPR screens in 12 cell lines. In addition, the average logs fold change is shown for 3 pan-essential genes and for the top 8 AML-biased dependencies nominated from the screens. **(B)** The list of genes showing top co-dependencies of UBE2T across 769 cell lines from Project Achilles (20Q3). The genes that belong to Fanconi anemia pathway are highlighted in bold.

**Figure S2.**
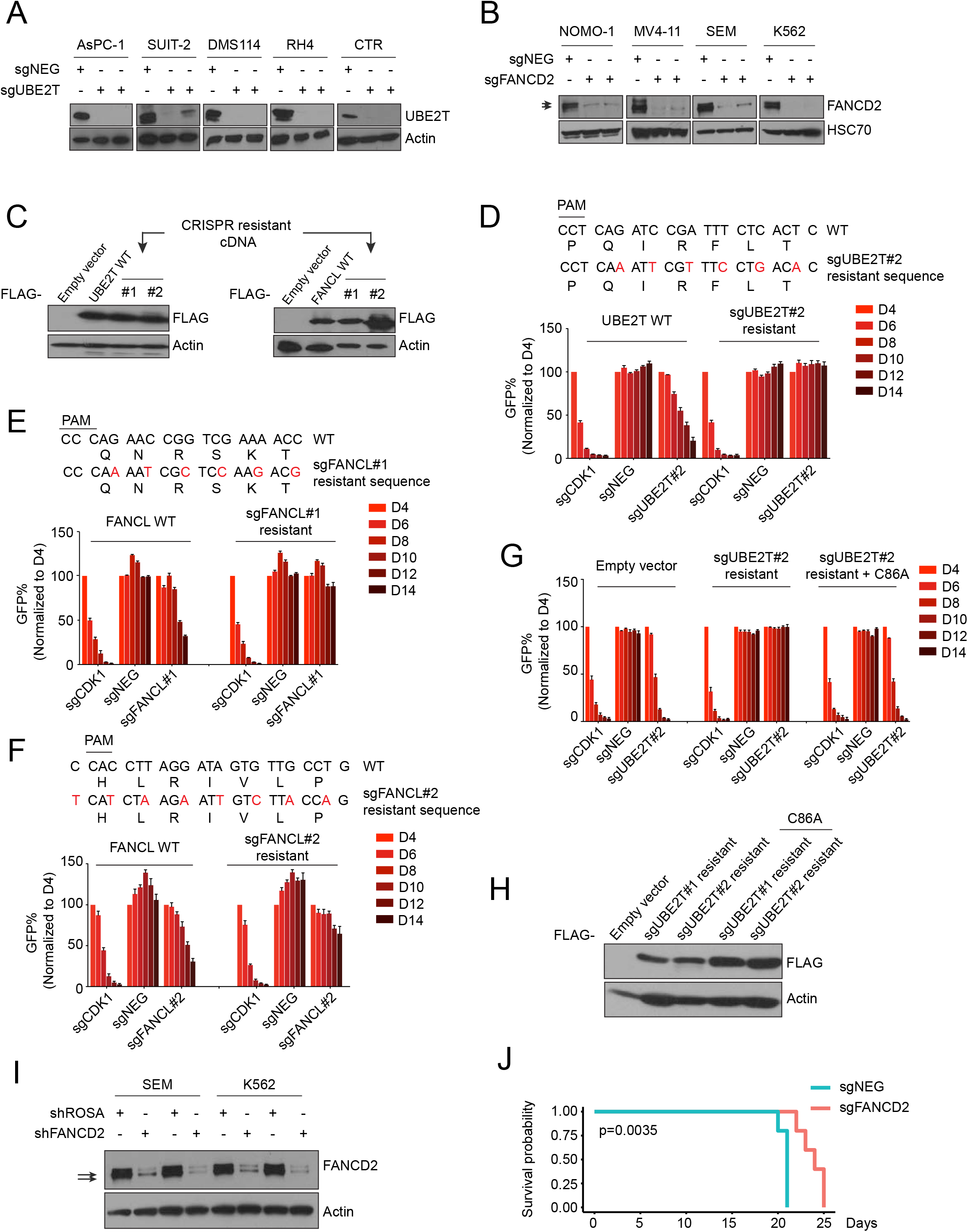
Validation of FA dependencies in AML maintenance. Related to Figure 2. **(A-B)** Western blot performed on whole cell lysates of the indicated cell lines following lentiviral transduction with indicated sgRNAs. **(C)** Western blot analysis in MOLM-13 cells of FLAG-tagged UBE2T (left) or FANCL (right) with wild-type sequence or harboring silent mutations of the sgRNA recognition site. **(D-G)** Competition-based proliferation assay in MOLM-13 cells expressing the indicated cDNAs (wild or CRISPR-resistant, labeled at the top) transduced with GFP-linked sgRNAs (labeled at the bottom). (n=3). **(H)** Western blot analysis of FLAG-tagged UBE2T in MOLM-13 cells, carrying wild-type or various mutations. **(I)** Western blot analysis following transduction with shRNAs targeting FANCD2. **(J)** Kaplan-Meier survival curves of NSG recipient mice transplanted with MOLM-13 cell infected with indicated sgRNA. Each sgRNA group contains 5 mice. A log-rank (Mantel-Cox) statistical test was used to calculate the p-value. All bar graphs represent the mean ± SEM.

**Figure S3.**
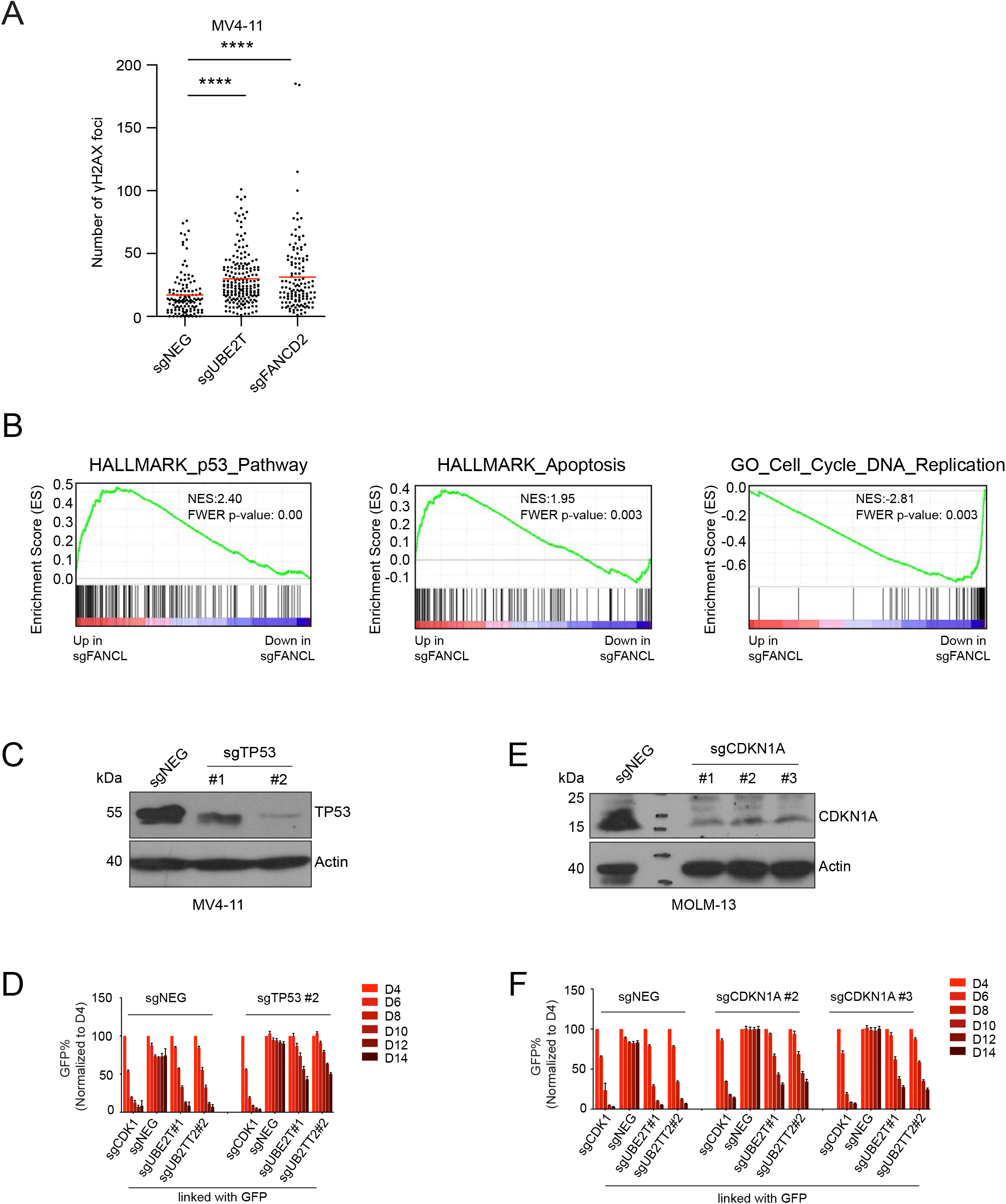
Targeting of FA genes leads to p53-dependent cell cycle arrest and cell death. Related to Figure 3. **(A)** The numbers of phospho-H2AX foci in MV-4-11-Cas9 cells infected with the indicated sgRNAs. Data were combined from three independent experiments. (sgROSA n=121, sgUBE2T n=184, and sgFANCD2 n=129). Statistical analysis was performed using one-way ANOVA followed by Dunnett’s multiple comparison test (**** p < 0.0001; ns, not significant). **(B)** Gene set enrichment analysis (GSEA) of RNA-seq data obtained from MOLM-13 cells lentivirally transduced with the indicated sgRNAs (Subramanian et al., 2005). Normalized enrichment score (NES) and family-wise error rate (FWER) *p*-value are shown. **(C)** Western blot performed on whole cell lysates prepared from MV4-11 cells on day 6 following transduction with the indicated sgRNAs. **(D)** Competition-based proliferation assays in MV4-11 cells following sequential sgRNA transduction. Negative sgRNA or TP53 sgRNAs were infected first, selected with neomycin, followed by transduction with the sgRNAs indicated at the bottom of the graph (linked with GFP). n=3. **(E)** Western blot analysis of whole cell lysates prepared from MOLM-13 cells on day 6 following transduction with the indicated sgRNAs. **(F)** Competition-based proliferation assays in MOLM-13 cells following sequential sgRNA transduction. Negative sgRNA or CDKN1A sgRNAs were infected first, selected with neomycin, followed by transduction with the sgRNAs indicated at the bottom of the graph (linked with GFP). n=3.

**Figure S4.**
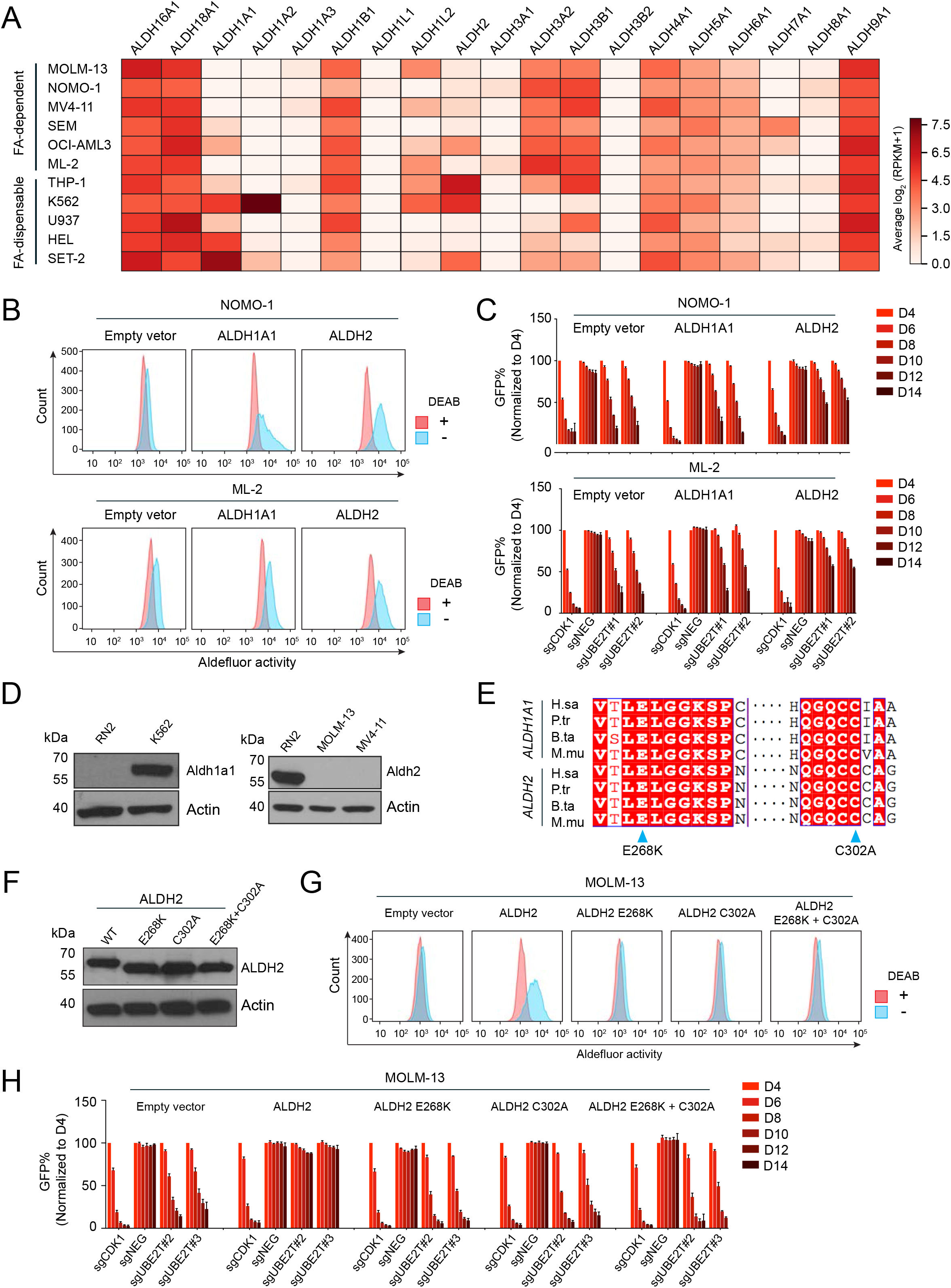
Additional experiments evaluating connection between ALDH genes and FA dependency in AML. Related to Figure 4. **(A)** RNA-seq analysis of mRNA levels for 19 ALDH genes in 11 leukemia cell lines. **(B)** Aldefluor analysis of NOMO-1 and MV4-11 cells transduced with FLAG-ALDH1A1 or FLAG-ALDH2. Lines were treated with 3.75 μM BAAA, with or without the ALDH inhibitor DEAB (15 μM). **(C)** Competition-based proliferation assay in Cas9+ NOMO-1 and ML-2 cells expressing the indicated cDNAs, followed by transduction with GFP-linked sgRNA. (n=3). **(D)** Western blot analysis of ALDH1A1 or ALDH2 proteins in the indicated leukemia cell lysates. **(E)** Multiple sequence alignment of ALDH1A1 and ALDH2. The catalytic residues are highlighted by blue triangles. **(F)** Western blot analysis of wild-type and catalytic mutants FLAG-ALDH2 in MOLM-13 lysates. **(G)** Aldefluor analysis of MOLM-13 cells transduced with FLAG-ALDH1A1 or FLAG-ALDH2. Lines were treated with 3.75 μM BAAA, with or without the ALDH inhibitor DEAB (15 μM). **(H)** Competition-based proliferation assay in MOLM-13 cells expressing wild-type or catalytic mutant ALDH2 using the indicated sgRNAs linked to GFP.

**Figure S5.**
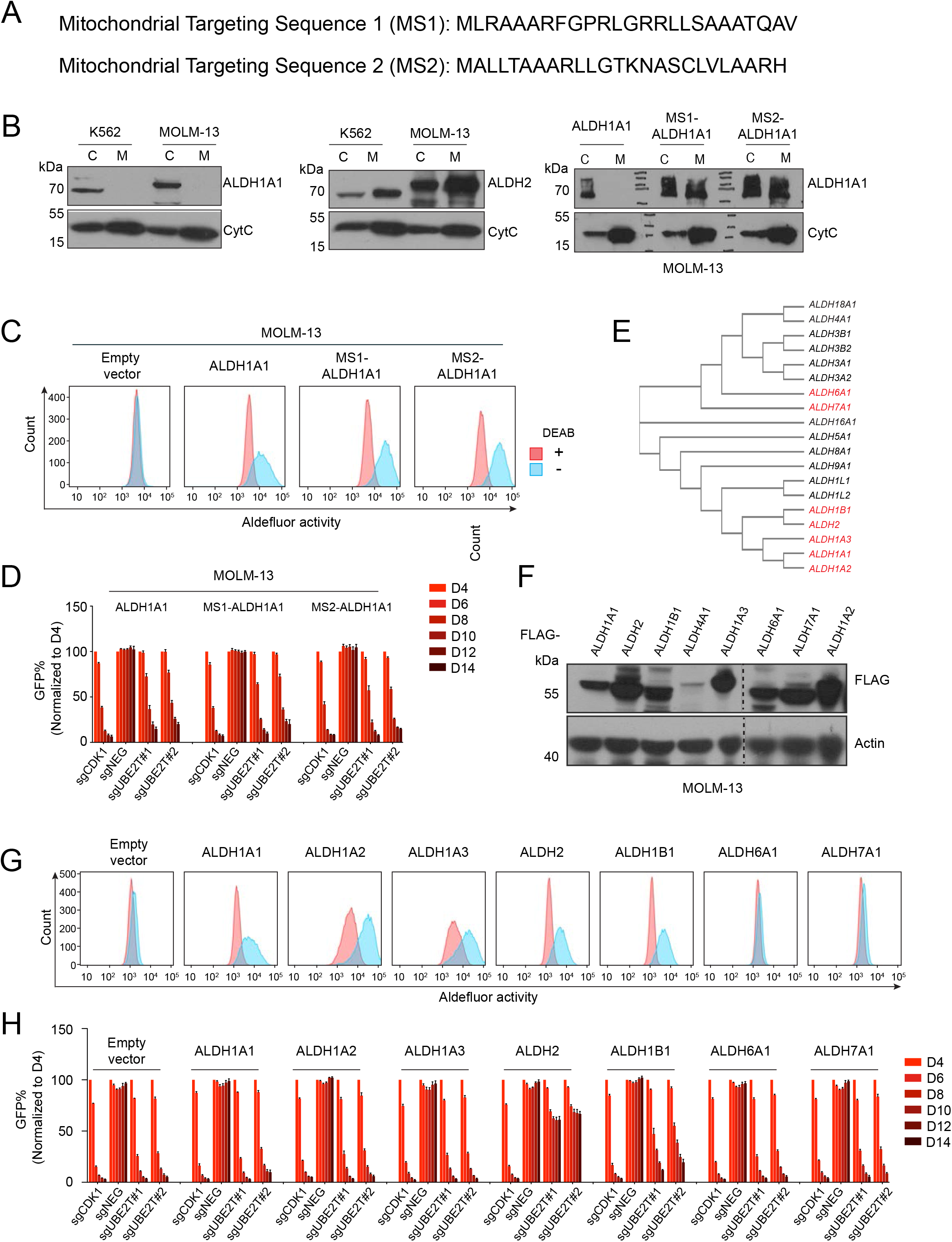
ALDH2 is the only ALDH gene that show synthetic lethality with Fanconi anemia complex. Related to Figure 4. **(A)** Two sequences of mitochondrial localization signal peptides fused to ALDH1A1. **(B)** Western blot analysis of cytoplasm (C) or mitochondrial (M) fractions of indicated cell lines. **(C)** Aldefluor analysis of MOLM-13 cells transduced with FLAG-ALDH1A1 or fusions with mitochondrial localization signal peptides (MS1 and MS2). Cells were treated with 3.75 μM BAAA, with or without the ALDH inhibitor DEAB (15 μM). **(D)** Competition-based proliferation assay in MOLM-13 cells expressing wild-type or mitochondrial localization signal peptide fused ALDH1A1 using the indicated sgRNAs linked to GFP. **(E)** Phylogenetic analysis of ALDH gene family. **(F)** Western blot analysis of whole cell lysates prepared from MOLM-13 cells transduced with the indicated FLAG-ALDH cDNAs. **(G)** Aldefluor assay of MOLM-13 cells transduced with the indicated ALDH genes. Lines were treated with 3.75 μM BAAA, with or without the ALDH inhibitor DEAB (15 μM). **(H)** Competition-based proliferation assay in MOLM-13 cells expressing the indicated ALDH genes. All bar graphs represent the mean ± SEM (n=3).

**Figure S6.**
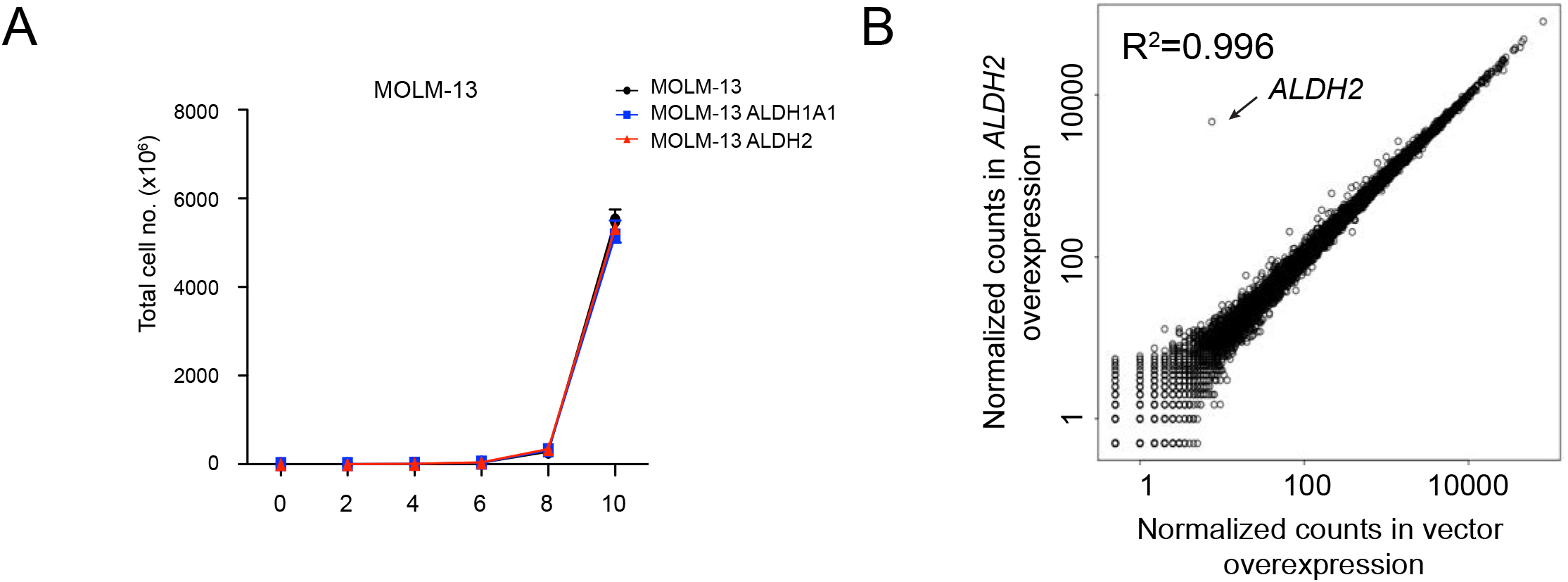
Re-expression of ALDH2 in MOLM-13 leads to minimal changes in cell proliferation and transcription. **(A)** Cell proliferation *in vitro* over the course of 10 days of the indicated cell lines. (n=3). **(B)** RNA-seq scatter plot analysis comparing fold-change of mRNA levels following ALDH2 overexpression versus empty vector in MOLM-13 cells.

**Figure S7.**
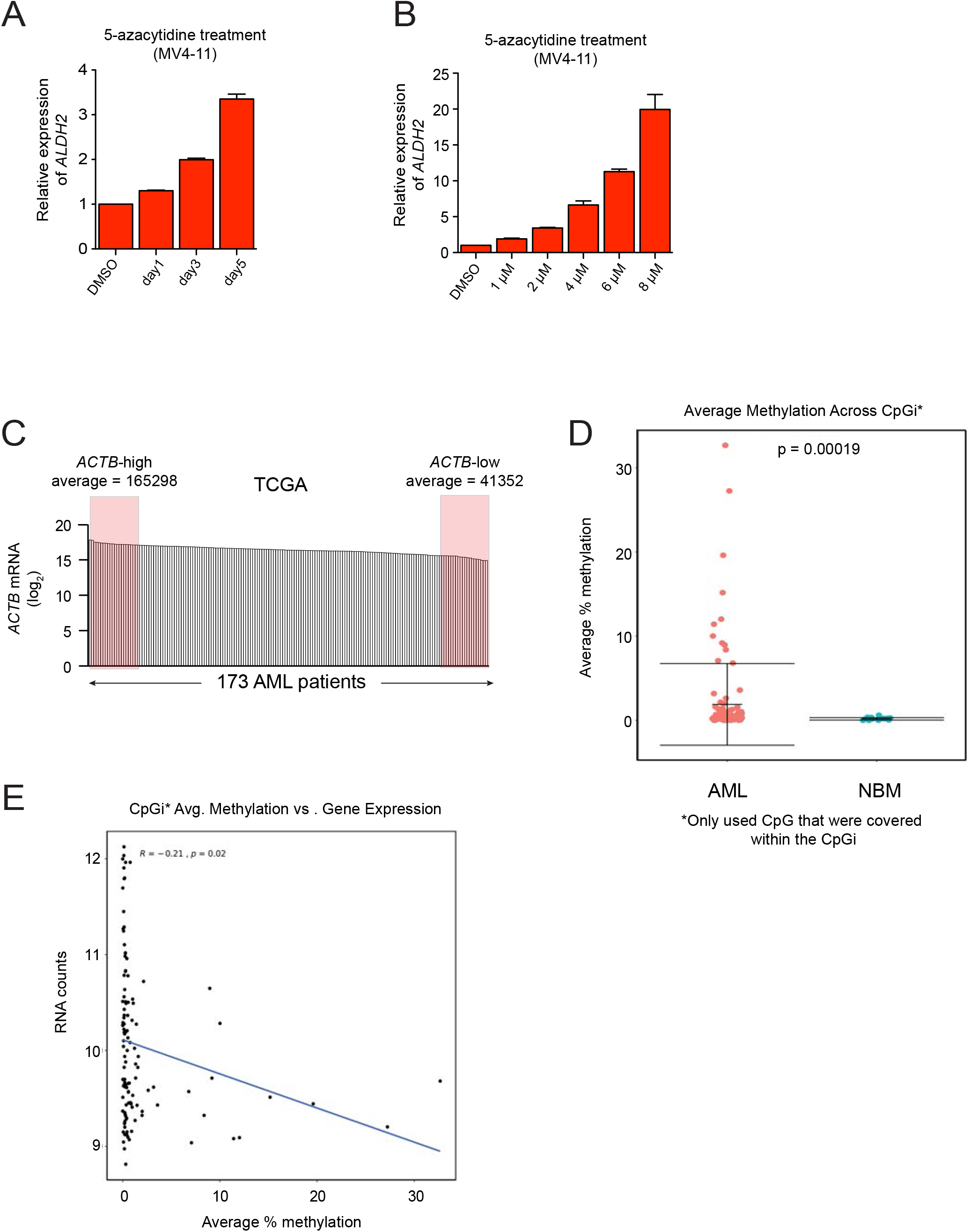
ALDH2 expression is silenced by aberrant DNA methylation in AML. Related to Figure 5. **(A-B)** RT-qPCR analysis of ALDH2 mRNA levels in MV4-11 cells following treatment with 5-azacytidine. In (A), 1 μM concentration was used. In (B), a 36 hour timepoint was used. **(C)** ACTB mRNA levels in 173 AM patient samples from TCGA. Shown are the samples classified as ACTB-high and ACTB-low. **(D)** Barplot of the average percent methylation of the CpGi. Each dot is representative of one patient. p-value from two tailed Student’s t-test. **(E)** Scatterplot of the average percent methylation of the CpGi versus the RNA expression level of ALDH2. Each dot is representative of one patient. Line of best fit was fit using a linear model. Correlation was calculated using Pearson’s method in R.

## REFERENCES

Akalin, A., Kormaksson, M., Li, S., Garrett-Bakelman, F.E., Figueroa, M.E., Melnick, A., and Mason, C.E. (2012). methylKit: a comprehensive R package for the analysis of genome-wide DNA methylation profiles. Genome Biol 13, R87.

Alcón, P., Shakeel, S., Chen, Z.A., Rappsilber, J., Patel, K.J., and Passmore, L.A. (2020). FANCD2– FANCI is a clamp stabilized on DNA by monoubiquitination of FANCD2 during DNA repair. Nat Struct Mol Biol 27, 240–248.

Alpi, A.F., Pace, P.E., Babu, M.M., and Patel, K.J. (2008). Mechanistic Insight into Site-Restricted Monoubiquitination of FANCD2 by Ube2t, FANCL, and FANCI. Molecular Cell 32, 767–777.

Barretina, J., Caponigro, G., Stransky, N., Venkatesan, K., Margolin, A.A., Kim, S., Wilson, C.J., Lehár, J., Kryukov, G.V., Sonkin, D., et al. (2012). The Cancer Cell Line Encyclopedia enables predictive modelling of anticancer drug sensitivity. Nature 483, 603–607.

Baylin, S.B., and Jones, P.A. (2016). Epigenetic Determinants of Cancer. Cold Spring Harb Perspect Biol 8, a019505.

Bryant, H.E., Schultz, N., Thomas, H.D., Parker, K.M., Flower, D., Lopez, E., Kyle, S., Meuth, M., Curtin, N.J., and Helleday, T. (2005). Specific killing of BRCA2-deficient tumours with inhibitors of poly(ADP-ribose) polymerase. 434, 6.

Ceccaldi, R., Sarangi, P., and D’Andrea, A.D. (2016). The Fanconi anaemia pathway: new players and new functions. Nat Rev Mol Cell Biol 17, 337–349.

Cereda, M., Mourikis, T.P., and Ciccarelli, F.D. (2016). Genetic Redundancy, Functional Compensation, and Cancer Vulnerability. Trends in Cancer 2, 160–162.

Chen, C.-H., Ferreira, J.C.B., Gross, E.R., and Mochly-Rosen, D. (2014). Targeting Aldehyde Dehydrogenase 2: New Therapeutic Opportunities. Physiological Reviews 94, 1–34.

Chen, Y., Koppaka, V., Thompson, D.C., and Vasiliou, V. (2012). Focus on Molecules: ALDH1A1: From lens and corneal crystallin to stem cell marker. Experimental Eye Research 102, 105–106.

Cremona, C.A., and Behrens, A. (2014). ATM signalling and cancer. Oncogene 33, 3351–3360.

Czerwińska, J., Poznański, J., Dębski, J., Bukowy, Z., Bohr, V.A., Tudek, B., and Speina, E. (2014). Catalytic activities of Werner protein are affected by adduction with 4-hydroxy-2-nonenal. Nucleic Acids Research 42, 11119–11135.

Dempster, J.M., Rossen, J., Kazachkova, M., Pan, J., Kugener, G., Root, D.E., and Tsherniak, A. (2019). Extracting Biological Insights from the Project Achilles Genome-Scale CRISPR Screens in Cancer Cell Lines (Cancer Biology).

Dobin, A., Davis, C.A., Schlesinger, F., Drenkow, J., Zaleski, C., Jha, S., Batut, P., Chaisson, M., and Gingeras, T.R. (2013). STAR: ultrafast universal RNA-seq aligner. Bioinformatics 29, 15–21.

Fan, X., Molotkov, A., Manabe, S.-I., Donmoyer, C.M., Deltour, L., Foglio, M.H., Cuenca, A.E., Blaner, W.S., Lipton, S.A., and Duester, G. (2003). Targeted Disruption of Aldh1a1 (Raldh1) Provides Evidence for a Complex Mechanism of Retinoic Acid Synthesis in the Developing Retina. MCB 23, 4637–4648.

Farmer, H., McCabe, N., Lord, C.J., Tutt, A.N.J., Johnson, D.A., Richardson, T.B., Santarosa, M., Dillon, K.J., Hickson, I., Knights, C., et al. (2005). Targeting the DNA repair defect in BRCA mutant cells as a therapeutic strategy. Nature 434, 917–921.

Feng, J., Liu, T., Qin, B., Zhang, Y., and Liu, X.S. (2012). Identifying ChIP-seq enrichment using MACS. Nat Protoc 7, 1728–1740.

Fernandez, E., Koek, W., Ran, Q., Gerhardt, G.A., France, C.P., and Strong, R. (2006). Monoamine Metabolism and Behavioral Responses to Ethanol in Mitochondrial Aldehyde Dehydrogenase Knockout Mice. Alcoholism Clin Exp Res 30, 1650–1658.

Garaycoechea, J.I., Crossan, G.P., Langevin, F., Daly, M., Arends, M.J., and Patel, K.J. (2012). Genotoxic consequences of endogenous aldehydes on mouse haematopoietic stem cell function. Nature 489, 571–575.

Garaycoechea, J.I., Crossan, G.P., Langevin, F., Mulderrig, L., Louzada, S., Yang, F., Guilbaud, G., Park, N., Roerink, S., Nik-Zainal, S., et al. (2018). Alcohol and endogenous aldehydes damage chromosomes and mutate stem cells. Nature 553, 171–177.

Gasparetto, M., Pei, S., Minhajuddin, M., Khan, N., Pollyea, D.A., Myers, J.R., Ashton, J.M., Becker, M.W., Vasiliou, V., Humphries, K.R., et al. (2017). Targeted therapy for a subset of acute myeloid leukemias that lack expression of aldehyde dehydrogenase 1A1. Haematologica 102, 1054–1065.

Gerber, J.M., Smith, B.D., Ngwang, B., Zhang, H., Vala, M.S., Morsberger, L., Galkin, S., Collector, M.I., Perkins, B., Levis, M.J., et al. (2012). A clinically relevant population of leukemic CD34ϩCD38Ϫ cells in acute myeloid leukemia. 119, 8.

Ghandi, M., Huang, F.W., Jané-Valbuena, J., Kryukov, G.V., Lo, C.C., McDonald, E.R., Barretina, J., Gelfand, E.T., Bielski, C.M., Li, H., et al. (2019). Next-generation characterization of the Cancer Cell Line Encyclopedia. Nature 569, 503–508.

Gilpatrick, T., Lee, I., Graham, J.E., Raimondeau, E., Bowen, R., Heron, A., Sedlazeck, F.J., and Timp, W. (2019). Targeted Nanopore Sequencing with Cas9 for studies of methylation, structural variants, and mutations (Genomics).

Glass, J.L., Hassane, D., Wouters, B.J., Kunimoto, H., Avellino, R., Garrett-Bakelman, F.E., Guryanova, O.A., Bowman, R., Redlich, S., Intlekofer, A.M., et al. (2017). Epigenetic Identity in AML Depends on Disruption of Nonpromoter Regulatory Elements and Is Affected by Antagonistic Effects of Mutations in Epigenetic Modifiers. Cancer Discov 7, 868–883.

Grabbe, C., Husnjak, K., and Dikic, I. (2011). The spatial and temporal organization of ubiquitin networks. Nat Rev Mol Cell Biol 12, 295–307.

Guo, J.-M., Liu, A.-J., Zang, P., Dong, W.-Z., Ying, L., Wang, W., Xu, P., Song, X.-R., Cai, J., Zhang, S.-Q., et al. (2013). ALDH2 protects against stroke by clearing 4-HNE. Cell Res 23, 915–930.

Hatakeyama, S. (2011). TRIM proteins and cancer. Nat Rev Cancer 11, 792–804.

Helleday, T., Petermann, E., Lundin, C., Hodgson, B., and Sharma, R.A. (2008). DNA repair pathways as targets for cancer therapy. Nat Rev Cancer 8, 193–204.

Hira, A., Yabe, H., Yoshida, K., Okuno, Y., Shiraishi, Y., Chiba, K., Tanaka, H., Miyano, S., Nakamura, J., Kojima, S., et al. (2013). Variant ALDH2 is associated with accelerated progression of bone marrow failure in Japanese Fanconi anemia patients. Blood 122, 3206–3209.

Hoang, V.T., Buss, E.C., Wang, W., Hoffmann, I., Raffel, S., Zepeda-Moreno, A., Baran, N., Wuchter, P., Eckstein, V., Trumpp, A., et al. (2015). The rarity of ALDH ^+^ cells is the key to separation of normal versus leukemia stem cells by ALDH activity in AML patients: Aldehyde dehydrogenase in acute myeloid leukemia. Int. J. Cancer 137, 525–536.

Hodges, E., Molaro, A., Dos Santos, C.O., Thekkat, P., Song, Q., Uren, P.J., Park, J., Butler, J., Rafii, S., McCombie, W.R., et al. (2011). Directional DNA Methylation Changes and Complex Intermediate States Accompany Lineage Specificity in the Adult Hematopoietic Compartment. Molecular Cell 44, 17–28.

Hoeller, D., and Dikic, I. (2009). Targeting the ubiquitin system in cancer therapy. Nature 458, 438–444.

Innan, H., and Kondrashov, F. (2010). The evolution of gene duplications: classifying and distinguishing between models. Nat Rev Genet 11, 97–108.

Jones, P.A., and Baylin, S.B. (2007). The Epigenomics of Cancer. Cell 128, 683–692.

Kaelin, W.G. (2005). The Concept of Synthetic Lethality in the Context of Anticancer Therapy. Nat Rev Cancer 5, 689–698.

Kafri, R., Springer, M., and Pilpel, Y. (2009). Genetic Redundancy: New Tricks for Old Genes. Cell 136, 389–392.

Kent, W.J., Sugnet, C.W., Furey, T.S., Roskin, K.M., Pringle, T.H., Zahler, A.M., and Haussler, D. The Human Genome Browser at UCSC. 11.

Kitagawa, K., Kawamoto, T., Kunugita, N., Tsukiyama, T., Okamoto, K., Yoshida, A., Nakayama, K., and Nakayama, K. (2000). Aldehyde dehydrogenase (ALDH) 2 associates with oxidation of methoxyacetaldehyde; in vitro analysis with liver subcellular fraction derived from human and *Aldh2* gene targeting mouse. FEBS Letters 476, 306–311.

Komander, D., and Rape, M. (2012). The Ubiquitin Code. Annu. Rev. Biochem. 81, 203–229.

Kottemann, M.C., and Smogorzewska, A. (2013). Fanconi anaemia and the repair of Watson and Crick DNA crosslinks. Nature 493, 356–363.

Kronke, J., Udeshi, N.D., Narla, A., Grauman, P., Hurst, S.N., McConkey, M., Svinkina, T., Heckl, D., Comer, E., Li, X., et al. (2014). Lenalidomide Causes Selective Degradation of IKZF1 and IKZF3 in Multiple Myeloma Cells. Science 343, 301–305.

Langevin, F., Crossan, G.P., Rosado, I.V., Arends, M.J., and Patel, K.J. (2011). Fancd2 counteracts the toxic effects of naturally produced aldehydes in mice. Nature 475, 53–58.

Li, H. (2018). Minimap2: pairwise alignment for nucleotide sequences. Bioinformatics 34, 3094–3100.

Li, H., Handsaker, B., Wysoker, A., Fennell, T., Ruan, J., Homer, N., Marth, G., Abecasis, G., Durbin, R., and 1000 Genome Project Data Processing Subgroup (2009). The Sequence Alignment/Map format and SAMtools. Bioinformatics 25, 2078–2079.

Love, M.I., Huber, W., and Anders, S. (2014). Moderated estimation of fold change and dispersion for RNA-seq data with DESeq2. Genome Biol 15, 550.

Lu, G., Middleton, R.E., Sun, H., Naniong, M., Ott, C.J., Mitsiades, C.S., Wong, K.-K., Bradner, J.E., and Kaelin, W.G. (2014). The Myeloma Drug Lenalidomide Promotes the Cereblon-Dependent Destruction of Ikaros Proteins. Science 343, 305–309.

Machida, Y.J., Machida, Y., Chen, Y., Gurtan, A.M., Kupfer, G.M., D’Andrea, A.D., and Dutta, A. (2006). UBE2T Is the E2 in the Fanconi Anemia Pathway and Undergoes Negative Autoregulation. Molecular Cell 23, 589–596.

Meyers, R.M., Bryan, J.G., McFarland, J.M., Weir, B.A., Sizemore, A.E., Xu, H., Dharia, N.V., Montgomery, P.G., Cowley, G.S., Pantel, S., et al. (2017). Computational correction of copy number effect improves specificity of CRISPR–Cas9 essentiality screens in cancer cells. Nat Genet 49, 1779–1784.

Nakahata, K., Uehara, S., Nishikawa, S., Kawatsu, M., Zenitani, M., Oue, T., and Okuyama, H. (2015). Aldehyde Dehydrogenase 1 (ALDH1) Is a Potential Marker for Cancer Stem Cells in Embryonal Rhabdomyosarcoma. PLoS ONE 10, e0125454.

Nene, A., Chen, C.-H., Disatnik, M.-H., Cruz, L., and Mochly-Rosen, D. (2017). Aldehyde dehydrogenase 2 activation and coevolution of its ∊PKC-mediated phosphorylation sites. J Biomed Sci 24, 3.

O’Connor, M.J. (2015). Targeting the DNA Damage Response in Cancer. Molecular Cell 60, 547–560.

Ohsawa, I., Nishimaki, K., Murakami, Y., Suzuki, Y., Ishikawa, M., and Ohta, S. (2008). Age-Dependent Neurodegeneration Accompanying Memory Loss in Transgenic Mice Defective in Mitochondrial Aldehyde Dehydrogenase 2 Activity. Journal of Neuroscience 28, 6239–6249.

Quinlan, A.R., and Hall, I.M. (2010). BEDTools: a flexible suite of utilities for comparing genomic features. Bioinformatics 26, 841–842.

Santos, M.A., Faryabi, R.B., Ergen, A.V., Day, A.M., Malhowski, A., Canela, A., Onozawa, M., Lee, J.-E., Callen, E., Gutierrez-Martinez, P., et al. (2014). DNA-damage-induced differentiation of leukaemic cells as an anti-cancer barrier. Nature 514, 107–111.

Schuurhuis, G.J., Meel, M.H., Wouters, F., Min, L.A., Terwijn, M., de Jonge, N.A., Kelder, A., Snel, A.N., Zweegman, S., Ossenkoppele, G.J., et al. (2013). Normal Hematopoietic Stem Cells within the AML Bone Marrow Have a Distinct and Higher ALDH Activity Level than Co-Existing Leukemic Stem Cells. PLoS ONE 8, e78897.

Senft, D., Qi, J., and Ronai, Z.A. (2018). Ubiquitin ligases in oncogenic transformation and cancer therapy. Nat Rev Cancer 18, 69–88.

Shi, J., Wang, E., Milazzo, J.P., Wang, Z., Kinney, J.B., and Vakoc, C.R. (2015). Discovery of cancer drug targets by CRISPR-Cas9 screening of protein domains. Nat Biotechnol 33, 661–667.

Simpson, J.T., Workman, R.E., Zuzarte, P.C., David, M., Dursi, L.J., and Timp, W. (2017). Detecting DNA cytosine methylation using nanopore sequencing. Nat Methods 14, 407–410.

Skaar, J.R., Pagan, J.K., and Pagano, M. (2014). SCF ubiquitin ligase-targeted therapies. Nat Rev Drug Discov 13, 889–903.

Sproul, D., Kitchen, R.R., Nestor, C.E., Dixon, J.M., Sims, A.H., Harrison, D.J., Ramsahoye, B.H., and Meehan, R.R. (2012). Tissue of origin determines cancer-associated CpG island promoter hypermethylation patterns. Genome Biol 13, R84.

Struhl, K. (2014). Is DNA methylation of tumour suppressor genes epigenetic? ELife 3, e02475.

Subramanian, A., Tamayo, P., Mootha, V.K., Mukherjee, S., Ebert, B.L., Gillette, M.A., Paulovich, A., Pomeroy, S.L., Golub, T.R., Lander, E.S., et al. (2005). Gene set enrichment analysis: A knowledge-based approach for interpreting genome-wide expression profiles. Proceedings of the National Academy of Sciences 102, 15545–15550.

Sullivan, K.D., Padilla-Just, N., Henry, R.E., Porter, C.C., Kim, J., Tentler, J.J., Eckhardt, S.G., Tan, A.C., DeGregori, J., and Espinosa, J.M. (2012). ATM and MET kinases are synthetic lethal with nongenotoxic activation of p53. Nat Chem Biol 8, 646–654.

The Cancer Genome Atlas Research Network (2013). Genomic and Epigenomic Landscapes of Adult De Novo Acute Myeloid Leukemia. N Engl J Med 368, 2059–2074.

Tomita, H., Tanaka, K., Tanaka, T., and Hara, A. (2016). Aldehyde dehydrogenase 1A1 in stem cells and cancer. Oncotarget 7, 11018–11032.

Verhaak, R.G.W., Wouters, B.J., Erpelinck, C.A.J., Abbas, S., Beverloo, H.B., Lugthart, S., Lowenberg, B., Delwel, R., and Valk, P.J.M. (2009). Prediction of molecular subtypes in acute myeloid leukemia based on gene expression profiling. Haematologica 94, 131–134.

Voulgaridou, G.-P., Anestopoulos, I., Franco, R., Panayiotidis, M.I., and Pappa, A. (2011). DNA damage induced by endogenous aldehydes: Current state of knowledge. Mutation Research/Fundamental and Molecular Mechanisms of Mutagenesis 711, 13–27.

Wang, A.T., and Smogorzewska, A. (2015). SnapShot: Fanconi Anemia and Associated Proteins. Cell 160, 354–354.e1.

Wang, R., Wang, S., Dhar, A., Peralta, C., and Pavletich, N.P. (2020). DNA clamp function of the monoubiquitinated Fanconi anaemia ID complex. Nature 580, 278–282.

Wei, J., Wunderlich, M., Fox, C., Alvarez, S., Cigudosa, J.C., Wilhelm, J.S., Zheng, Y., Cancelas, J.A., Gu, Y., Jansen, M., et al. (2008). Microenvironment Determines Lineage Fate in a Human Model of MLL-AF9 Leukemia. Cancer Cell 13, 483–495.

Zuber, J., McJunkin, K., Fellmann, C., Dow, L.E., Taylor, M.J., Hannon, G.J., and Lowe, S.W. (2011). Toolkit for evaluating genes required for proliferation and survival using tetracycline-regulated RNAi. Nat Biotechnol 29, 79–83.

